# Exaptation and de novo mutations transcend cryptic variations as drivers of adaptation in yeast

**DOI:** 10.1101/2024.03.26.586634

**Authors:** Shreya Routh, Richard J. Lindsay, Ivana Gudelj, Riddhiman Dhar

## Abstract

Many organisms live in predictable environments with periodic variation in growth condition which can allow populations to accumulate cryptic genetic variations. Cryptic variations can facilitate adaptation to new environments, as observed in evolution experiments with a ribozyme and a protein. Whether the same holds for cell populations remains unclear. Alternatively, living in a near-constant condition can lead to loss of nonessential cellular functions, which could be maladaptive in new environments. Through laboratory evolution experiments in yeast, we show that populations grown in a predictable nutrient-rich environment for 1000 generations start to lose their ability to respond and adapt to new stressful environments. Growth of yeast populations in the nutrient-rich environment was associated with modest fitness increase in this environment, metabolic remodeling, and increased lipid accumulation. In novel stressful environments, however, these populations generally had reduced fitness, except in salt-stress where lipid accumulation seemed to provide osmotic protection. We further found that adaptation to stressors was primarily driven by de novo mutations, with very little contribution from the mutations accumulated prior to the exposure to stressors. Thus, our work suggests that in the absence of occurrence of new environments, natural populations might not accumulate cryptic variations that could be beneficial for adaptation to these environments. In addition, presence of selection in predictable condition in natural populations may purge away some of the cryptic variations. Taken together, these findings raise questions about persistence of cryptic variations in natural populations and their importance in evolutionary adaptation.

## Introduction

Many organisms live in predictable environments with periodic variation in growth condition, for instance, variation in nutrient abundance, or in temperature, such as the one associated with day-night cycles. On the one hand, living in a predictable environment can enable populations to explore the genotype space and accumulate cryptic genetic variations. Cryptic variations may have little or no benefit in that environment, but can help in adaptation to new environments (Rouzic and Carlborg 2008; Paaby and Rockman 2014), as shown by evolution experiments with a ribozyme (Hayden et al. 2011) and a fluorescent protein (Zheng et al. 2019). Further, cryptic genetic variations can influence epistatic interactions, and thereby, the evolution of proteins towards novel functions (Baier et al. 2019). However, it is unclear whether these observations can be extended to populations (McGuigan and Sgrò 2009; Paaby and Rockman 2014). Experiments in bacteria, that were limited to adaptation to new nutrients (Blount et al. 2012; Rigato and Fusco 2016; Izutsu and Lenski 2022), showed that standing genetic variations do not always facilitate adaptation to new nutrients (Izutsu and Lenski 2022). Further, experiments in yeast revealed an interplay between standing genetic variation and de novo mutations that determined evolution (Ament-Velásquez et al. 2022).

Alternatively, living in a predictable environment can optimize and streamline the cellular processes that help an organism thrive in that condition (D’Souza and Kost 2016). For instance, costly genes can be rapidly lost during experimental evolution when they are not required for the environment (Nilsson et al. 2005; Koskiniemi et al. 2012), such as biosynthetic genes whose product are already available (D’Souza and Kost 2016). Moreover, near-constant growth condition and a regular supply of essential nutrients are thought to be the reasons behind occurrence of fewer genes and smaller genomes in intracellular parasites compared to their free-living counterparts (Moran 2002; Sakharkar et al. 2004; McCutcheon and Moran 2011; Corradi 2015). While such functional loss could be beneficial in one environment (Koskiniemi et al. 2012), they might be detrimental in others.

To test how population fitness is shaped by cryptic genetic variations and loss of function during growth in a predictable environment, we experimentally evolved populations of the yeast *Saccharomyces cerevisiae* in a nutrient-rich medium for 1000 generations. All evolved lines increased fitness in this medium, and showed metabolic changes that increased respiratory activity, utilization of the glyoxylate cycle, and lipid accumulation compared to the ancestors. However, apart from an increased fitness under salt stress, likely due to lipid-mediated osmo-protection, the evolved lines showed lower fitness in stressful environments. Subsequent selection experiments revealed poor evolvability of these lines in new environments, including crashing of two populations in oxidative stress. Mutation analysis revealed accumulation of high frequency non-synonymous mutations in these lines, some of which led to loss of gene functions. Moreover, mutations accumulated in rich medium did not contribute to evolutionary adaptation to new environments, which was predominantly driven by de novo mutations. Taken together, these results illustrate loss of nonessential functions in a predictable growth environment, highlight the role of metabolic adaptation and de novo mutations in fitness gain, and reveal a lack of contribution from cryptic genetic variations towards adaptation.

## Results

### Fitness of yeast populations increases during 1000 generations in one environment

To test for emergence of cryptic variations and loss-of-function mutations in a non-stressful environment, we experimentally evolved three replicate subpopulations (A lines), drawn from a single population of *S. cerevisiae,* in one nutrient-rich environment for 1000 generations. In each round, we grew the ancestral lines in nutrient-rich YPD medium for 24 hours (∼10 generations) (Fig. 1A) and transferred one thousandth of the population to fresh medium. We repeated this process for 100 rounds for a total of ∼1000 generations (Fig. 1A) and referred to the evolved populations as ‘N’ (no external stressor) lines. During each round, cells went through a periodic and predictable variation in nutrient availability. We tracked cell densities at the end of each round (Fig. 1B), and the effective population size for the replicate N lines were 8.4 × 10^4^, 7.7 × 10^4^ and 8.8 × 10^4^ respectively.

**Figure 1.**
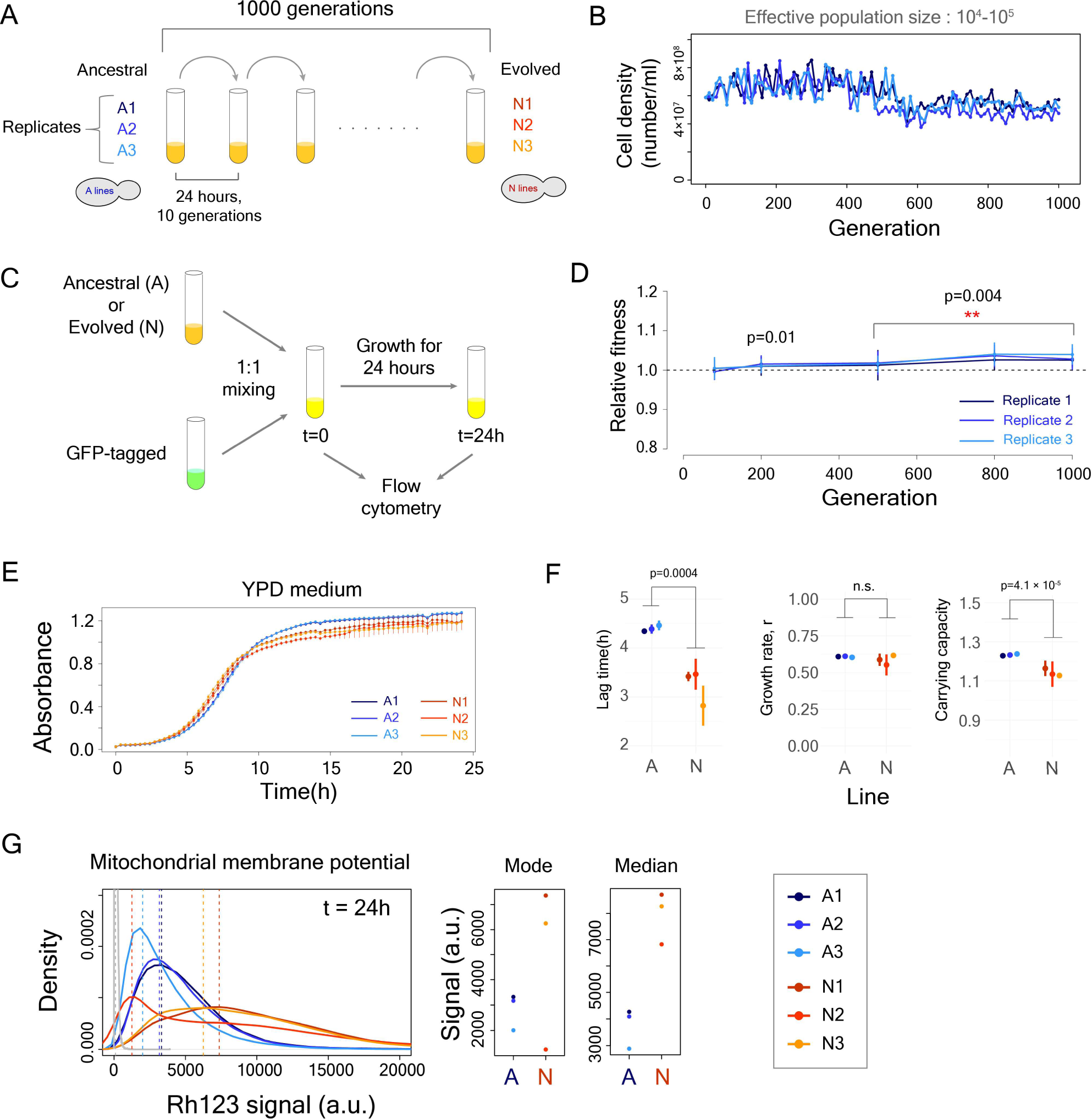
Yeast populations adapt to nutrient-rich YPD medium for 1000 generations. **(A)** A schematic diagram of laboratory evolution experiment in YPD medium. **(B)** Cell density (number/ml) at the end of every 24-hour cycle tracked through 1000 generations. **(C)** Schematic diagram of the competition assay to determine relative fitness of evolved lines compared to the ancestral lines (Three measurements for each line, and tested using one-sample Wilcoxon signed-rank test for all replicates from all lines). **(D)** Relative fitness of the evolved lines increased over the course of the evolution experiment. **(E)** Growth curve of the ancestral and evolved lines in YPD medium (in triplicates). **(F)** Estimates of lag time, growth rate and carrying capacity showed shorter lag and lower carrying capacity in the evolved lines compared to the ancestral lines (Wilcoxon rank-sum test for comparison between all replicates of ancestral and evolved lines) **(G)** Distribution of Rhodamine 123 signal in the ancestral and the evolved lines at 24 hours of growth. We compared the distribution of each evolved line against its corresponding ancestral line (Kolmogorov-Smirnov test, p<10^-10^ for each line). The distribution in grey is for unstained negative control and the dotted vertical lines show mode values of the distributions.

Earlier experiments with *Escherichia coli* and *S. cerevisiae* had revealed continued adaptation of populations to single environments for thousands of generations (Lenski et al. 2015; Johnson et al. 2021). Thus, we measured the fitness of the evolved lines (N1-N3) relative to the ancestors at different time points during evolution (Fig. 1C). We found that they all had increased fitness by generation 200, and showed ∼2.5% (± 1.8%), ∼2.8% (± 2.8%) and ∼4.0% (± 2.5%) increase in fitness, respectively, by generation 1000 (Fig. 1D, Wilcoxon signed-rank test for all measurements from all replicate lines).

Cells grown in a batch culture can increase fitness through reducing lag time or increasing growth rate or both (Gresham and Dunham 2014). Compared to the ancestors, all replicate N lines showed shortened lag time (∼26% reduction) and reduced carrying capacity (∼7% reduction), but did not show significant change in growth rate (Fig 1E-F). The shorter lag time in the N lines suggested that they could utilize nutrients quicker than the ancestors, which could lead to higher competitive fitness (Gemma et al. 2018) (Fig. 1D).

### The increase in fitness is associated with metabolic remodeling

Shortening of lag-phase suggested quicker processing of nutrients in the evolved lines, thereby enabling faster synthesis of the required biomolecules for initiating exponential growth. For a Crabtree-positive yeast like *S. cerevisiae*, glucose is processed through both fermentation and respiration (Pfeiffer and Morley 2014; Malina et al. 2021). Thus, faster processing of glucose requires higher expression or activity of enzymes involved in fermentation and the tri-carboxylic acid (TCA) cycle. Higher respiratory activity has been associated with shortening of lag time in yeast strains, albeit for shifting from one carbon source to another (Gemma et al. 2018; Vermeersch et al. 2019).

To test for changes in respiratory activity, we quantified mitochondrial membrane potential of the populations using the Rhodamine 123 (Rh123) stain (Fig. 1G; Fig. S1). The evolved lines showed significantly higher mitochondrial membrane potential compared to their ancestors (Kolmogorov-Smirnov test, p<10^-10^ for each line, Fig. 1G). Although the N2 line had a higher median signal than the ancestral lines, it showed a very heterogeneous population, including a subpopulation with lower signal than the ancestral lines, as shown by the mode (Fig. 1G). The increase in respiratory activity allowed evolved lines to grow faster in glycerol, a carbon source that required respiration (Fig. S2). Moreover, the evolved lines showed a substantially lower percentage of respiration deficient cells (<3%), that are spontaneously generated due to deficiency in mitochondrial function (Dimitrov et al. 2009), compared to the ancestors (∼10-15%) (Fig. S3).

In addition, yeast can utilize a shorter metabolic path - known as the glyoxylate shunt - in which the enzyme isocitrate lyase (ICL1) converts isocitrate to malate in just two reactions instead of five reactions in the full TCA cycle (Dduntze et al. 1969; Maaheimo et al. 2001; VillasBôas et al. 2005) (Fig. 2A), thereby possibly speeding up processing of glucose. Therefore, we measured expression levels of isocitrate lyase (ICL1) to see whether the evolved lines had changes in activity of the glyoxylate shunt (Fig. 2A), by introducing a C-terminal GFP tag and quantifying expression in two clones from each transformed line by flow cytometry. All evolved lines showed higher expression of ICL1 at early-log phase (4 hours), but showed lower expression at stationary phase (24 hours) compared to the ancestors (Fig. 2B; Kolmogorov-Smirnov test for comparison between clones of corresponding evolved and ancestral lines, p<10^-10^). We observed the opposite expression pattern for the pyruvate decarboxylase (*PDC1*) gene (Fig. S4).

**Figure 2.**
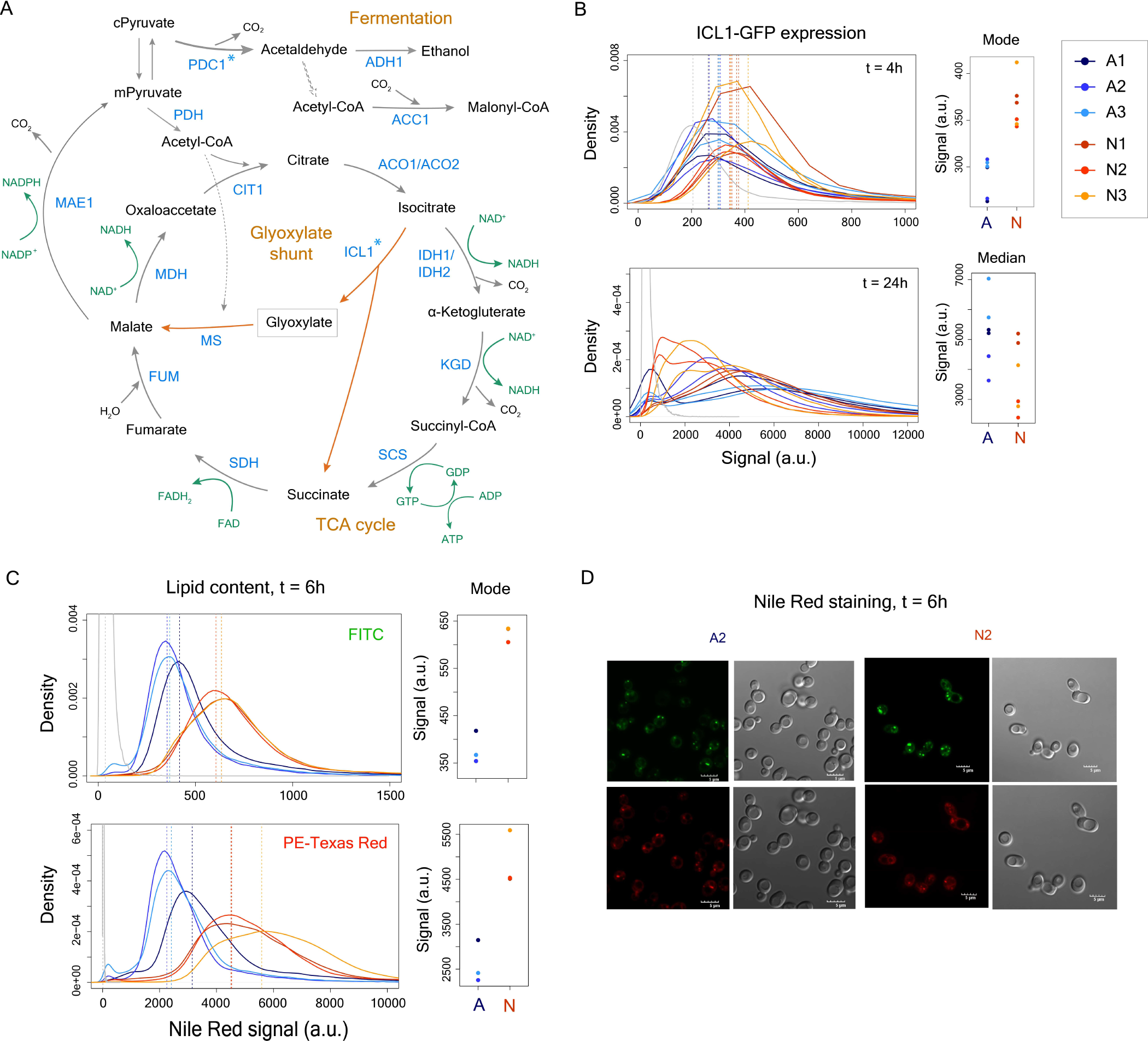
Metabolic remodeling leads to higher lipid accumulation in the evolved lines. **(A)** TCA cycle and glyoxylate shunt in *S. cerevisiae.* cPyruvate – cytosolic pyruvate, mPyruvate – mitochondrial pyruvate. **(B)** Expression of ICL1-GFP fusion protein measured by flow cytometry at 4 hours and 24 hours of growth (Kolmogorov-Smirnov test for comparison between clones of corresponding evolved and ancestral lines, p<10^-10^ at t=4h and at t=24h). Mode signal was estimated for unimodal distributions, whereas median signal was estimated for multi-modal distributions. **(C)** Lipid content measured at 6 hours of growth using Nile red staining (Kolmogorov-Smirnov test for comparison between corresponding evolved and ancestral lines, p<10^-10^ for all comparisons) **(D)** Microscopic images of cells from A2 and N2 lines stained with Nile red.

The glyoxylate shunt generates excess oxaloacetate molecules than the TCA cycle (Fig. 2A) that can be channelized towards citrate which has been associated with increased lipid production (Tang et al. 2015; Hu et al. 2019). We therefore quantified the lipid content in the evolved lines after 6 and 24 hours of growth through flow cytometry using the Nile red stain (Greenspan et al. 1985) (Fig. 2C,D). The evolved lines showed significantly higher signal than the ancestral lines in both FITC and PE-Texas Red channels at 6 hours (Fig. 2C) and at 24 hours (Fig. S5), suggesting increased lipid accumulation in these lines (Kolmogorov-Smirnov test, p<10^-10^ for all comparisons). These lipids appeared as droplets (Fig. 2D), which contain storage lipids consisting of triacyl glycerol and steryl esters (Klug and Daum 2014). We also observed an increase in cell size, estimated by forward scatter (FSC), in all evolved lines (Kolmogorov-Smirnov test, p<10^-10^ for all comparisons) (Fig. S6).

### Adaptation to a single environment leads to lower fitness and poor adaptability in new environments

To test for existence of cryptic genetic variations and loss-of-function mutations, we quantified fitness of the evolved lines in new environmental conditions. Accumulation of cryptic variations is likely to increase fitness in new environments or facilitate faster adaptation towards these environments, as has been observed in earlier experiments (Hayden et al. 2011; Zheng et al. 2019). We quantified growth of all lines in minimal growth medium (SCD), supplemented with salt stress (SCD+1M NaCl), oxidative stress (SCD + 4mM H_2_O_2_), or antifungals (SCD+0.08 µg/ml caspofungin and SCD+3 µg/ml flucytosine). In minimal medium, the evolved lines showed lower growth rate and carrying capacity (Fig. 3A,B). In addition, the N2 line showed >2-fold increase in lag-time compared to the ancestors (p<10^-4^, One-way ANOVA with Tukey HSD). In salt stress, however, the evolved lines had shortened lag-time, increased growth rate but lower carrying capacity (Fig. 3C,D). This was likely due to higher accumulation of intracellular lipids such as triacylglycerol (Klug and Daum 2014), which could act as an osmo-protectant (Hohman 2002) in the evolved lines.

**Figure 3.**
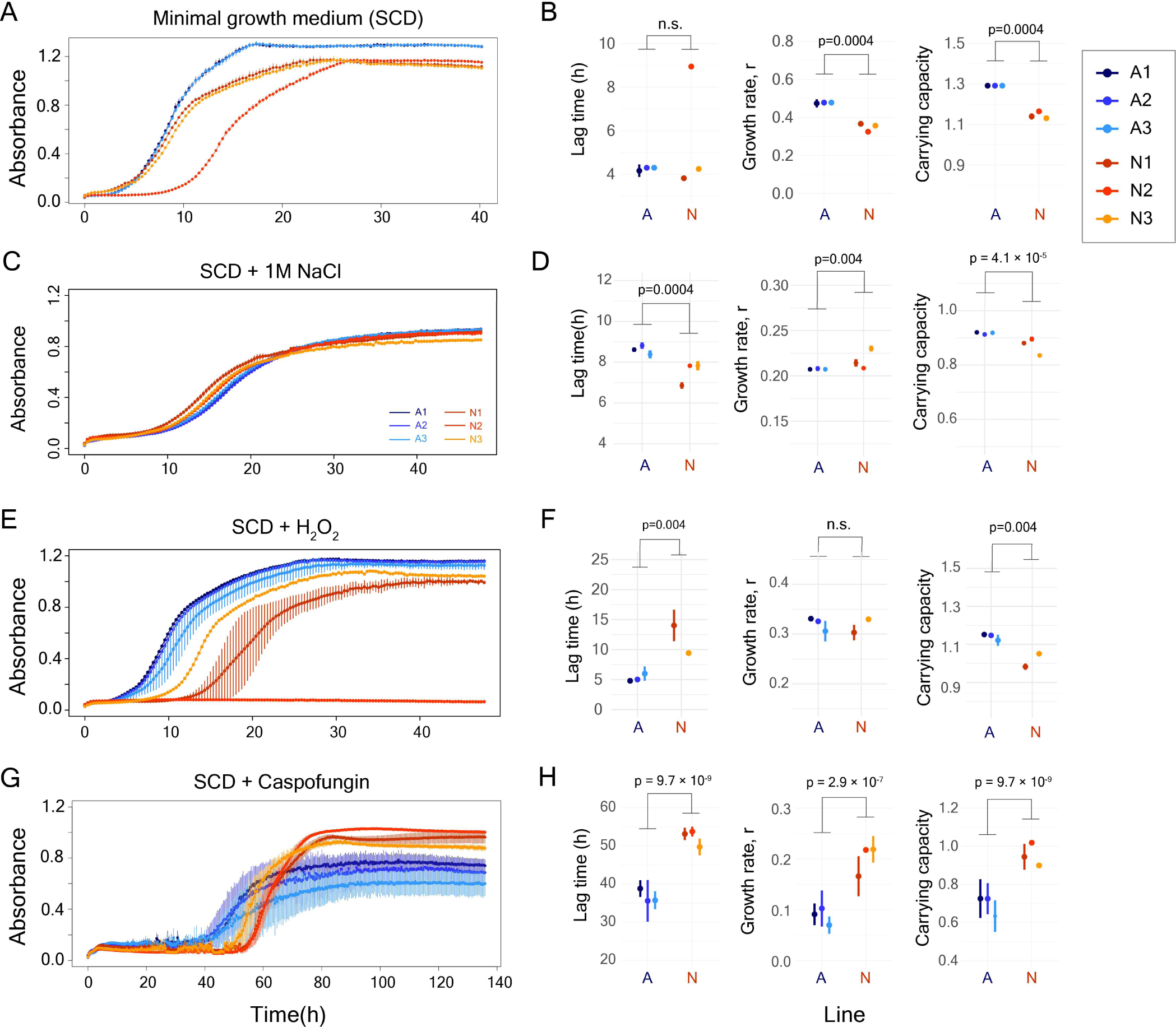
Evolved lines show lower fitness in new environmental conditions. **(A), (C), (E), (G)** – Growth curves of ancestral and evolved lines in SCD medium, SCD supplemented with 1M NaCl, 4mM H_2_O_2_, and 0.08 µg/ml caspofungin, respectively. **(B), (D), (F) (H)–** Estimation of lag time, growth rate and carrying capacity for ancestral and evolved lines in SCD medium, SCD supplemented with 1M NaCl, 4mM H_2_O_2_, and 0.08 µg/ml caspofungin, respectively. Wilcoxon rank-sum test was performed for comparisons of lag-time, growth rate and carrying capacity between evolved and ancestral lines.

In contrast, N2 line did not grow in oxidative stress, while N1 and N3 lines had increased lag-time, suggesting they had lower competitive fitness than the ancestral lines (Fig. 3E,F). Lower fitness of the evolved lines in oxidative stress was associated with changes in the expression of the gene *SOD1* (Fig. S7), that encodes for a superoxide dismutase and is a key component of oxidative stress response (Tsang et al. 2014). In antifungal compounds, the evolved lines had increased lag-time (Fig. 3G,H and Fig. S8), again suggesting lower competitive fitness of the evolved lines in these conditions. Taken together, these results showed that prolonged growth in one predictable condition had lowered the capabilities of the evolved lines to tolerate stressful environments in general, except osmotic stress.

Next, we tested whether the evolved lines had an advantage in adapting to stressful environments by performing a selection experiment with all A and N lines in salt stress, oxidative stress, and antifungal compounds. Briefly, we grew populations for 48 hours in each condition before transferring ∼10^5^ cells to fresh medium with the same stressor. This process was repeated for 10 rounds. We tracked cell densities by absorbance at the end of each round and measured growth properties after 10 rounds (Fig. 4; Fig. S9-S12). We limited this experiment to 10 rounds to evaluate the role of cryptic variations in adaptation and to minimize the impact of de novo mutations that are likely to arise in a longer selection experiment (Ament-Velásquez et al. 2022).

**Figure 4.**
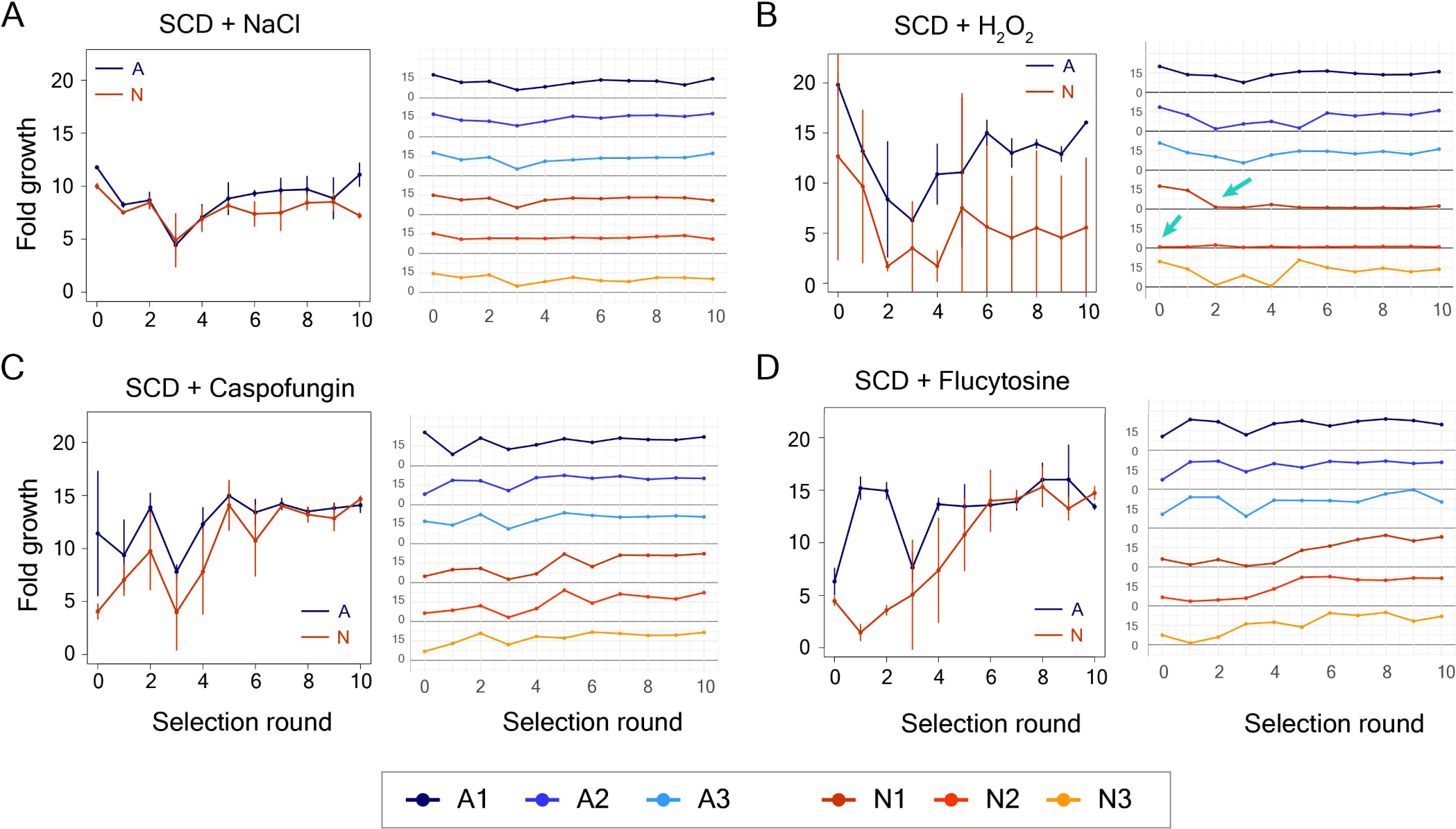
Evolved lines do not show better adaptability to new environments. Growth of A and N lines over 10 selection rounds, with each round lasting for 48 hours, in SCD + 1M NaCl **(A),** SCD + 4mM H_2_O_2_ **(B),** SCD + 0.08 µg/ml Caspofungin **(C)**, and SCD + 3 µg/ml flucytosine **(D)**. Plots on the left side show mean fold growth, along with ±1.0 standard deviation, of ancestral and evolved lines over 10 rounds of selection. The error bars show standard deviation values obtained from results of three replicate lines. Plots on the right show fold-growth in individual replicate lines over 10 rounds of selection.

In salt stress, we did not see much difference in the adaptation trajectories between the ancestral (A) and the evolved (N) lines (Fig. 4A). In oxidative stress, however, the line N2 did not grow (Fig. 4B), as expected from fitness measurements (Fig. 3E). Further, A1-3 and N3 showed a decrease in cell densities after the initial 4-5 rounds before recovering to similar densities by round 10, while N1 showed a gradual decrease in cell density by the end of round 3 that did not recover (Fig. 4B). In caspofungin and flucytosine, the N lines showed lower cell densities than the A lines for the first few rounds, but later recovered to the same cell densities as the ancestral lines (Fig. 4C,D). Growth curve analysis revealed that both A and N lines adapted to the antifungal compounds, but did not have significant difference in their growth profiles in the selection conditions (Fig. S10 and S12). Thus, the N lines generally had lower fitness than the A lines, both when initially exposed to stressors and when selected over 10 rounds. These results suggested that the N lines did not have increased adaptive capability to new environments.

### N lines have distinct mutations that hit several stress response genes

To map genomic changes that occurred during adaption in the nutrient-rich environment, we sequenced genomes of all A and N lines, and identified mutations that were unique to the N lines (Fig. 5A,B). The replicate lines showed distinct mutational signatures with a small fraction of mutations being shared between the replicate lines (Fig. 5A; File S1). We observed 175, 48 and 124 mutations in N1, N2 and N3 lines respectively, of which 55, 42 and 37 mutations showed population frequency of at least 5.0% (Fig. 5C). High frequency single nucleotide polymorphisms (SNPs), present in at least 5.0% frequency in the population, caused 17, 13 and 14 nonsynonymous changes (Fig. 5D). In addition, copy number variation analysis revealed only one segmental amplification in the N2 line (Fig. S13), as well as a mtDNA copy number increase in the N3 line (Fig. S14).

**Figure 5.**
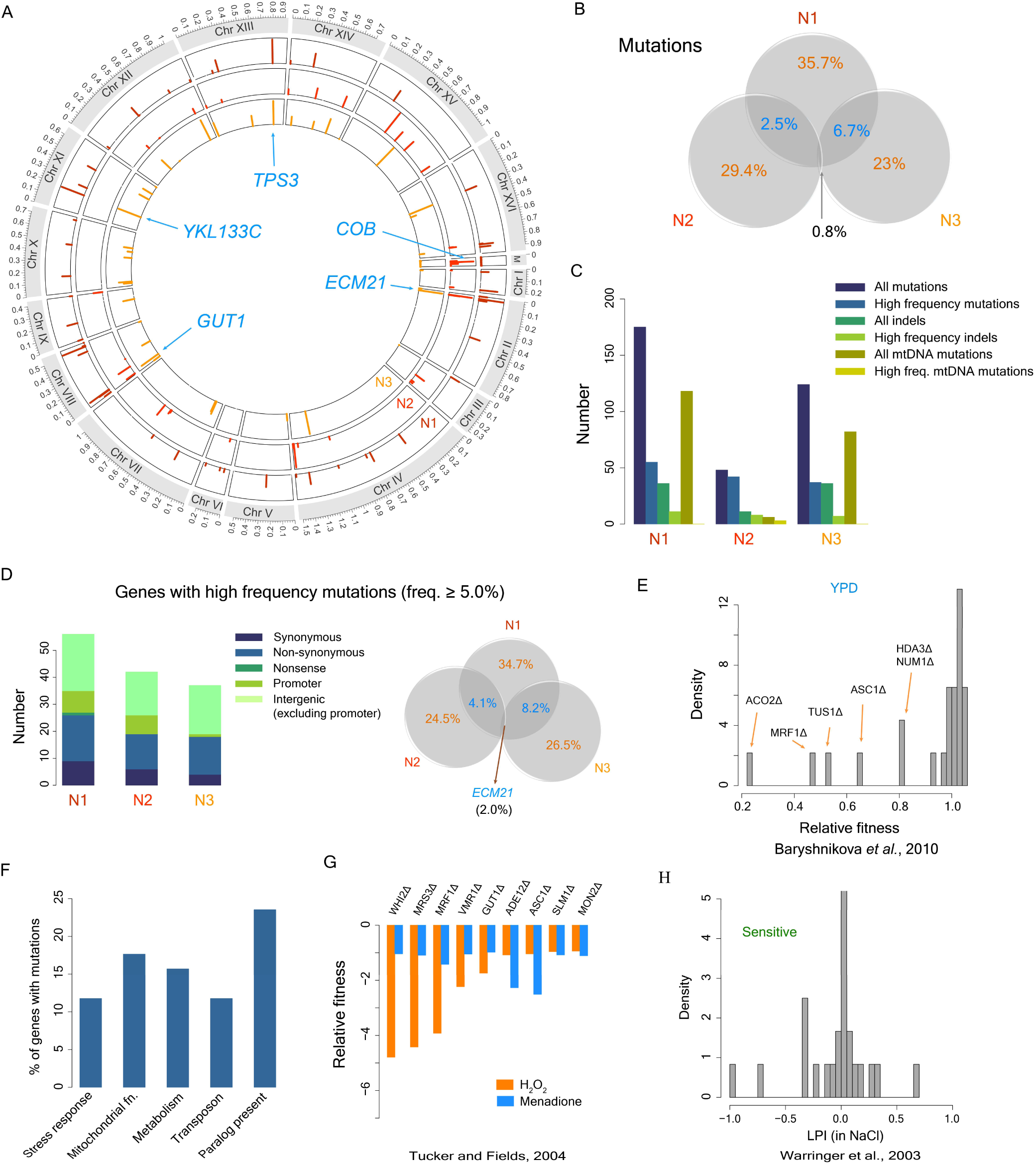
Replicate N lines show distinct mutational patterns. **(A)** Circular bar plot showing mutations in the genomes of the replicate lines N1, N2 and N3 that are not present in ancestral lines. Bar heights show population frequency of the mutations. **(B)** Venn diagram showing common and unique mutations in the replicate lines. **(C)** Bar plot showing number of all mutations, high frequency mutations (population frequency ≥ 5.0 %), number of all indels, high frequency indels, all mtDNA mutations, and high frequency mtDNA mutations in the evolved lines. **(D)** Bar plot showing number of synonymous, nonsynonymous, nonsense, promoter, and intergenic mutations. Venn diagram on the right shows the percentage of unique and common genes with mutations in the evolved lines. **(E)** Distribution of fitness relative to wild-type strain in YPD (data from Baryshnikova *et al*., 2010) of individual knock-outs of genes that contained nonsynonymous mutations in our evolved lines. **(F)** Percentage of genes containing mutations that were associated with stress-response, mitochondrial function, metabolism, transp

One gene, *ECM21/ART2*, contained SNPs as well as frameshift indels in all replicate lines (Fig. 5A,D), and was therefore, non-functional. *ECM21* is associated with endocytosis of amino acid transporters (Lussier et al. 1997), and is induced in response to excess amino acid, starvation, and other stress (Lin et al. 2008; Ivashov et al. 2020). Interestingly, earlier experiments with repeated cycles of selection for growth and starvation, and lysine limited growth conditions have also observed occurrences of nonsense and frameshift mutations in the *ECM21* gene (Wloch-Salamon et al. 2017; Hart et al. 2021).

In addition, the N1 line harbored a nonsense mutation in the *KRE6* gene (W526 to Stop) that was present at ∼54% frequency in the population. *KRE6* is a non-essential gene in the yeast *S. cerevisiae*, and is involved in β-1,6 glucan biosynthesis, although its exact enzymatic activity is unknown (Roemer and Bussey 1991; Kurita et al. 2011). The N2 line contained a high frequency (∼91%) nonsynonymous mutation in the cytochrome B (*COB*) gene, present in the mtDNA (Fig. S15) and encoding for a subunit of the complex III of the electron transport chain (Wenz et al. 2007). This mutation could also have an impact on mitochondrial function (Fig. 1G) and on oxidative stress response in the N2 line (Fig. 3E).

To check for the most possible deleterious impact of nonsynonymous mutations in the evolved lines, we analyzed the fitness of the corresponding gene knockouts in rich medium using data from Baryshnikova et al. (2010). Majority of the gene knockouts (>80%) had fitness values close to 1 in rich medium, suggesting minimal impact of mutations, even with potentially complete loss of function of these genes, on fitness in this medium (Fig. 5E).

Among the 51 genes containing high-frequency mutations, ∼13% of the genes were associated with stress response and ∼17% of the genes were associated with mitochondrial function (Fig. 5F). Very interestingly, individual knock-outs of nine of these genes were earlier found to make cell populations highly sensitive to oxidative stress (Tucker and Fields 2004) (Fig. 5G). Among these, mutations in five genes (*MRS3, MRF1, SLM1, VMR1,* and *ADE12*) were observed only in the N1 line, mutation in one gene (*WHI2*) was observed only in the N2 line, and mutations in two genes (*MON2* and *ASC1*) were observed only in the N3 line (File S1). Among these mutations, the M1I mutation in the *WHI2* gene, that was present in the N2 line at a frequency of ∼95%, led to a loss of start codon, and therefore, a likely loss of function. The gene *WHI2* is involved in activation of general stress response (Kaida et al. 2002). As knocking out *WHI2* gene was earlier found to make cells sensitive to oxidative stress (Tucker and Fields 2004), this mutation was likely to be a reason behind the extreme sensitivity of the N2 line to oxidative stress.

Similarly, knockouts of 10 genes containing mutations in N lines were earlier reported to be sensitive to salt stress (Fig. 5H; LPI<0) (Warringer et al. 2003). One stress-response associated gene, *HKR1* that encodes for an osmo-sensor in the HOG signaling pathway (Yabe et al. 1996), contained a nonsynonymous L346P mutation at a frequency of ∼91% in the N3 line. Despite these mutations, the N lines did not show lower fitness in salt than the ancestral lines, suggesting that the increased lipid accumulation in N lines could compensate for any detrimental effect of these mutations.

### De novo mutations drive adaptation to new environments

Next, we asked whether the mutations accumulated in the nutrient-rich medium had any role in the subsequent adaptation of the N lines to the environmental stressors. To address this question, we sequenced the genomes of all replicate A and N lines at the end of 10 rounds of selection in the antifungal compounds, caspofungin and flucytosine, as all lines showed significant adaptation to these two environments.

If a mutation accumulated in the nutrient-rich medium facilitated adaptation to environmental stressors, its frequency would rise after selection (Fig. 6A), and the magnitude of the frequency increase is likely to determine its importance in adaptation. Conversely, if a mutation had a deleterious impact on fitness in the stressor, its frequency would decrease and eventually it could be lost from a selected population (Fig. 6A). We found several mutations in the N lines whose frequency decreased or mutations which were lost after caspofungin (Fig. 6A,B,C) or flucytosine selection (Fig. S16A,B,C), but we did not find any mutation that was present in the N lines and showed a large increase in frequency after selection (File S3, S4). These results suggested that the mutations accumulated in the nutrient-rich medium in the N lines had neutral or deleterious fitness effect in the selection conditions, and had very little role in adaptation.

**Figure 6.**
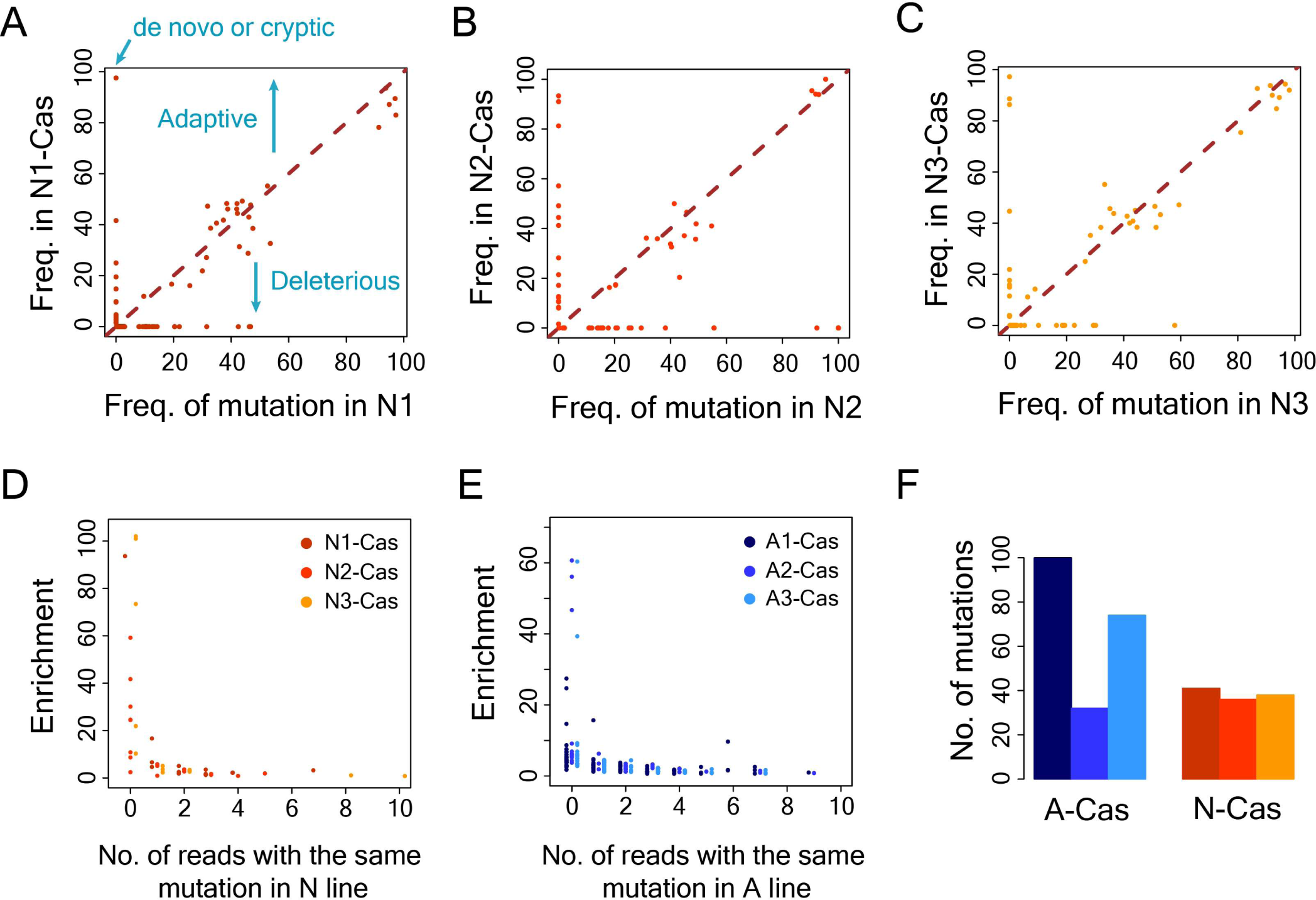
Adaptation to caspofungin is driven by de novo mutations. **(A)** Frequency of mutations in the N1 line before (N1) and after (N1-Cas) caspofungin selection **(B)** Frequency of mutations in the N2 line before (N2) and after (N2-Cas) caspofungin selection **(C)** Frequency of mutations in the N1 line before (N3) and after (N3-Cas) caspofungin selection **(D)** The y-axis shows enrichment of mutations in the caspofungin-selected N lines and the x-axis shows the number of nonduplicate reads in the genomic data of the corresponding N line in which the mutation was detected **(E)** The y-axis shows enrichment of mutations in the caspofungin-selected A lines and the x-axis shows the number of nonduplicate reads in the genomic data of the corresponding A line in which the mutation was detected **(F)** Distribution of number of mutations in caspofungin-selected A and N lines (denoted by A-Cas and N-Cas, respectively). oson, and had a paralog present. **(G)** Fitness of gene knock-outs relative to wild-type strain in medium containing H_2_O_2_ or menadione (Tucker and Fields, 2004) of knock-outs of genes that harbored nonsynonymous mutations in N lines. **(H)** Fitness of gene knock-outs relative to wild-type strain in salt (Warringer et al. 2003).

We observed several mutations that rose to high frequency after selection in caspofungin and flucytosine, but were not detected in the N lines ((Fig. 6A,B,C; Fig. S16A,B,C; File S3, S4). It is possible that these mutations represented cryptic variations, but were below the mutation detection threshold in our pipeline (Fig. 6A). Therefore, we tested whether these mutations appeared in even one read in the genomic data of the corresponding N line, which could then indicate these mutations to be of cryptic origin.

Further, for each mutation, we calculated an enrichment score which considered the frequency of the mutation in the selected line and in the corresponding N line, as well as the sequencing depth in these lines (see methods). High enrichment suggested a substantial increase in frequency of a mutation in the selected lines over the N lines after accounting for variation in sequencing depth. However, our analysis showed that all mutations showing high enrichment (>20) in the selected lines were not detected even in a single read in the N lines (Fig. 6D; Fig. S16D). This suggested that the mutations that rose to high frequency in the selected lines, and therefore primary drivers of adaptation in these conditions, were likely to be de novo mutations. We observed similar patterns in the selected lines that arose from the ancestral A lines, which did not undergo adaptation to nutrient-rich environment in our experiment (Fig. 6E, Fig. S16E). Further, we did not see any systemic differences in the number of mutations among the selected lines arising from A lines and N lines (Fig. 6F, Fig. S16F). These results collectively suggested that adaptation to environmental stressors was primarily driven by de novo mutations, and not cryptic genetic variations.

## Discussion

Taken together, our work shows that prolonged growth in a predictable environment can lead to loss of nonessential functions, and can degrade ability of populations to adapt to new environments. These findings contrast with earlier studies (Hayden et al. 2011; Zheng et al. 2019) which showed that cryptic variations accumulated during stabilizing selection could facilitate adaptation to new environments. Similarly, an earlier study in *E. coli* also reached the same conclusion, although new environments in their experiment were limited to new carbon sources (Rigato and Fusco 2016). There are several possible reasons for this difference. First, earlier experiments maintained a stabilizing selection regime where fitness did not change prior to the exposure to new environments. Stabilizing selection has been suggested to result in accumulation of cryptic genetic variations (Waddington 1957). In comparison, yeast populations in our experiment showed an increase in fitness, although modest, in the nutrient rich condition. Positive selection in this condition could purge away some of the cryptic variations. A second possible reason could be the difference in the mutation rates between the current study and the earlier work. Earlier studies used elevated mutation rates in the form of PCR mutagenesis (in the case of in vitro studies) (Hayden et al. 2011; Zheng et al. 2019) or chemical mutagenesis (with *E. coli* populations) (Rigato and Fusco 2016). In addition, these mutagenesis methods have well-known mutational biases (Sega 1984; D’Costa et al. 2023) which could also impact the results.

The evolved lines showed remarkably similar parallelism in metabolic remodeling that led to higher respiratory activity and increased lipid accumulation compared to the ancestors. The metabolic remodeling had a positive impact on fitness in two environments. Higher respiratory activity enabled the evolved lines to grow faster on carbon sources that required respiration. In addition, higher storage lipid accumulation, in the form of triacylglycerol, provided osmo-protection and helped evolved lines perform better in osmotic stress. These observations illustrate a role of exaptation in evolution, as metabolic remodeling occurred as an adaptation to growth in one environment but were beneficial for adaptation to other environments. Further, these results highlight how the effects of exaptation could also be misconstrued as the effects of cryptic genetic variations.

Although it is difficult to exactly pinpoint the impact of each mutation that accumulated in the rich medium on fitness, because of epistatic interactions with other mutations (Domingo et al. 2019; Johnson et al. 2023), it is likely that some of these mutations did not have any fitness effect and simply rose to high frequency by hitchhiking with beneficial mutations (Smith and Haigh 1974; Fay and Wu 2000). Therefore, it is surprising that none of them contributed to subsequent adaptation to new environments.

The findings of this study raise questions about the occurrence and maintenance of cryptic genetic variations in natural populations and their impact on evolutionary adaptation. Cryptic genetic variations were defined to explain the presence of mutations in a population that were beneficial for survival in new environments which a population had not yet seen (Dobzhansky 1941; Waddington 1957). However, the present work shows that in the prolonged absence of the new environments, capabilities to adapt to these conditions might degrade over time. Thus, appearance of new environments at specific time intervals might be necessary for maintaining these functions. Transient appearance of new environmental conditions can enable populations to acquire adaptive mutations that may persist over time as cryptic variations when the environment returns to normal condition. Thus, it might be necessary to carefully reexamine how cryptic variations arise and persist in natural populations (Paaby and Rockman 2014). Moreover, laboratory evolution experiments show that populations continue to adapt to an environment even after many thousands of generations (Lenski et al. 2015; Johnson et al. 2021). Presence of sustained selection in natural populations can remove some of the cryptic variations which might be beneficial for adaptation to new environments. Thus, the impact of cryptic variations in evolutionary adaptation of natural populations may not be as significant as originally thought.

## Materials and Methods

### Laboratory evolution experiment

*Saccharomyces cerevisiae* strain BY4741 was used for laboratory evolution experiment. The experiment was initiated from three biological replicates grown from a single yeast population in Yeast Peptone Dextrose (YPD) medium containing 1% (w/v) yeast extract, 2% (w/v) peptone and 2% (w/v) glucose. These replicate populations were referred to as A1, A2 and A3 lines. In each round, each replicate population was grown in 5ml YPD medium in a 50 ml glass tube for ∼24 hours in an incubator shaker maintained at 30°C and with agitation at 200 rpm. After 24 hours, the cultures reached stationary phase, and 5 µl of saturated culture was transferred to a fresh tube containing 5ml YPD medium. Each transfer round consisted of ∼10 generations (∼log_2_1000). This process was continued for 100 rounds and yeast lines were adapted for 1000 generations. The lines adapted for 1000 generations in YPD medium were referred to as N1, N2 and N3 lines. To keep track of growth of the cells during transfer experiments, Optical Density at 600 nm (OD_600_) was measured at the end of every growth cycle using a spectrophotometer.

To determine the relationship between OD_600_ and cell number, a plate count assay was implemented. Saturated yeast cultures were plated (80µl) at different dilutions (1000X, 5000X and 10000X) and each dilution in three replicates on YPD plates containing 1.5% agar. The plates were incubated for 24 hours at 30°C and the colonies were counted. The average number of cells per ml per OD at 600 nm was estimated as ∼6.64×10^6^, which was considered for calculating cell numbers in all further experiments. The effective population size was calculated for each replicate as a simple harmonic mean between the highest cell number and the lowest cell number over the evolution experiment.

### Competition assay

To assess for changes in fitness during the evolution experiment, a competition assay was performed for populations at generations 80, 200, 500, 800 and 1000, and relative fitness values with respect to the ancestors were quantified. For competition assay, a *S. cerevisiae* BY4741 strain with genome integrated GFP gene under the control of the RPL35A promoter at the site of *HIS3* gene was used as the reference strain. Cells of each line was grown in YPD medium for 24 hours, then was diluted (75-fold) and grown in fresh YPD for 24 hours, followed by a final dilution (50-fold) in YPD medium and 4 hours growth. At this stage, cells of each line were mixed with cells of the GFP-tagged strain in 1:1 ratio (1 ml each), pelleted and the supernatant was discarded. Cells were resuspended in 1 ml of Phosphate Buffered Saline (PBS, containing 137 mM NaCl, 10 mM Na_2_HPO_4_, and 2 mM KH_2_PO_4_, pH 7.4), and measurement from 100,000 cells were obtained by flow cytometer (BD LSR Fortessa Cell Analyzer) at the start of the competition assay (t = 0 h) (Fig. S17). The same cell mixture was diluted 1000-fold in fresh medium and was grown at 30°C and 200 rpm for ∼24 hours. After 24 hours of growth, 1 ml of sample was washed and resuspended in 1 ml PBS. The percentage of GFP-positive cells were then quantified using flow cytometry (Fig. S18).

Relative fitness of each of the ancestral and evolved lines was calculated with respect to the GFP-positive reference strain, to eventually estimate relative fitness of the evolved lines with respect to the ancestral lines. The percentages of GFP-positive and GFP-negative cells were obtained, and relative fitness (*w*) values were calculated following the method in Dhar et al. (2013). Briefly, following measures were calculated.

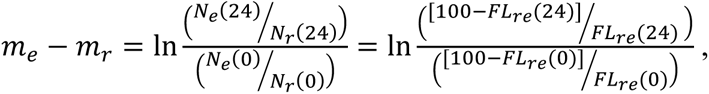

Where m_e_ was the Malthusian fitness of an evolved line, m_r_ was the Malthusian fitness of the reference strain, FL_re_(0) was the percentage of GFP-tagged reference strain at t=0, and FL_re_(24) was the percentage of GFP-tagged reference strain at t=24h.

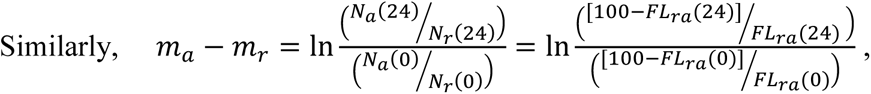

Where m_a_ was the Malthusian fitness of an evolved line, m_r_ was the Malthusian fitness of the reference strain, FL_ra_(0) was the percentage of GFP-tagged reference strain at t=0, and FL_ra_(24) was the percentage of GFP-tagged reference strain at t=24h.

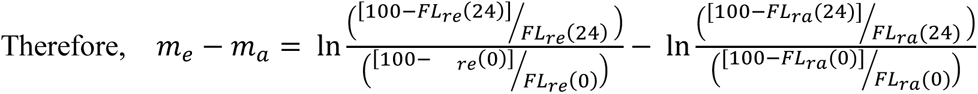

In our experiment, growth was limited to ∼1000 fold increase in each transfer, therefore, m_a_ was approximated as m_a_ = m_av_ = ln(1,000)/(1 day)= 6.9078 d^-1^. Selection coefficient *s* was defined as the difference between Darwinian fitness between an evolved line and the ancestral lines.

Thus, s = (m_e_ – m_a_)/m_a_ = w – 1, where s>0 or w>1 indicated a fitness advantage of the evolved line over the ancestral line (Lenski et al., 1991). Relative fitness of the evolved lines at generations 80, 200, 500, 800, and 1000 were estimated using this method.

### Analysis of growth curves

The growth parameters of A and N-lines in their native and new environments were determined through measuring growth curves in these environments. From overnight cultures of A and N-lines, ∼10^5^ cells from each culture were transferred to fresh medium in a 48 well plate to a total volume of 640 µl. The cells were grown at 30°C in Tecan Spark Multimode Microplate Reader and optical density at 600nm was measured every 20 mins for 48 hours. Growth curves were determined in nutrient-rich YPD medium, SCD (Synthetic Complete Defined, 0.67% YNB w/v + 0.079% complete synthetic supplement w/v + 2% w/v glucose), SCD + 0.08µg/ml caspofungin, SCD + 3µg/ml flucytosine, SCD + 1M NaCl, and SCD + 4mM H_2_O_2_. A stock solution of 1 mg/ml caspofungin was prepared in molecular grade water and filtered with 0.2 µm syringe filters. It was freshly diluted 10-fold and added to SCD to a working concentration of 0.08 µg/ml used in the experiments. Similarly, a stock solution of 15 mg/ml flucytosine was prepared with molecular grade water, filter sterilized, and freshly diluted 10-fold to working concentration of 3 µg/ml. For salt stress, 5.844 g of NaCl was added along with YNB, CSM and D-glucose to 100ml of solution and autoclaved. H_2_O_2,_ (3% v/v ≈ 900 mM) was freshly diluted in SCD to a working concentration of 4mM just before experiments. Lag time, growth rate and carrying capacity were estimated by fitting a logistic growth equation to the growth curve data using non-linear least squares method (‘nls’) in R (Fig. S19). Briefly, cell density was expressed as 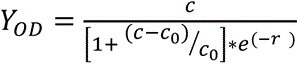, where Y_OD_ was the optical density of cells at 600nm at a time point *t*, c_0_ was the initial cell density (at t=0), c was the carrying capacity, and *r* was the growth rate. Lag time was determined from a growth curve as follows. Lines were fit to sliding windows of 20 data points, and their slope values were calculated. The maximum value of slope was identified and the latest time point at which the slope of the fitted line became lower than 0.01 compared to the maximum slope was chosen as the lag time. Wilcoxon rank-sum test was performed for comparison between all replicate measurements of all ancestral lines and replicate measurements of all evolved lines.

### Measurement of mitochondrial membrane potential

The mitochondrial membrane potential of the A and N lines was quantified using rhodamine123 staining. Cells from each line were grown overnight in 5ml YPD medium at 30°C, were diluted 75-fold in fresh medium, and were allowed to grow for another 24 hours. The cells reached stationary phase at the end of 24 hours, and 2 ml cells were collected for further processing. Cells were centrifuged and the supernatant was discarded. Cells were then washed twice with PBS (8000 rpm, 5 mins) and then stained with rhodamine 123 dye added at a final conc. of 10 μM (from 10 mM stock). The cells were incubated at 30 °C for 15 mins with gentle shaking inside an incubator. After 15 mins, cells were washed and resuspended in 1 ml PBS buffer. Flow cytometry was done to quantify mitochondrial membrane potential (Fig. S20). In addition, microscopic images were also captured for visualization of the mitochondria at 100X magnification under green filter in a confocal microscope (FLUOVIEW FV3000 confocal laser scanning microscope, Olympus).

### Quantification of respiration deficient cells

Percentage of respiration deficient cells in ancestral and evolved lines were determined by plating cells in synthetic complete medium containing 0.1% glucose and 3% glycerol v/v (SCDG). Briefly, cells were grown overnight in SCD medium at 30°C, then diluted in fresh medium and were grown for 4 hours. At this stage, cells were washed and resuspended in PBS, and 200 cells for each line were plated on SCDG medium. The plates were incubated at 30°C for 96 hours. Percentage of respiration deficient cells were determined by counting the number of small and large colonies in the SCDG plates. The assay was done in three technical replicates for each line.

### Construction of GFP-tagged fusion proteins

C-terminal GFP tagged fusion proteins of the genes *ICL1*, *PDC1* and *SOD1* were created following the protocol described by Longtine et al. 1998. Primers for tagging of these genes were taken from Howson et al., 2005, where F2 and R1 primers for each gene were obtained (Table S1). GFP-tag for each gene was amplified using these primers from a *S. cerevisiae* strain with PDR5-GFP construct. Amplification was done with 0.5 μl Q5 polymerase enzyme (NEB), using 10 μl Q5 buffer, 1 μl dNTP, 2.5 μl each of the forward and reverse primers, 32.5 μl molecular grade water and ∼1 ng of PDR5-GFP genomic DNA as template. The amplification protocol consisted of incubation at 94°C for 3 min, followed by 10 cycles of incubation at 94°C for 15s, at 50°C for 30 s, at 72°C for 2 min, followed by 15 cycles of incubation at 94°C for 15s, at 72°C for 2 min, and a final incubation at 72°C for 10 min. Length of the amplified products (2348 bp) was confirmed by 0.8% agarose gel electrophoresis and the products were transformed into A and N lines through homologous recombination with Histidine as the auxotrophic selection marker.

Chemically competent yeast cells from A and N lines were prepared by adding overnight grown cells (150 µl) to 25 ml of YPD and incubating at 30°C for 4-5 hours with occasional shaking. The cells were pelleted down and washed with 800 µl of 0.1 M lithium acetate. Cells were then resuspended in 50 µl of lithium acetate, and 5ul of salmon sperm DNA (ssDNA, boiled for 5 minutes at 95°C just before adding and kept on ice) along with 30 µl of PCR amplified product were added. The tubes were flicked and allowed to sit for 5 min. A PLI mixture (300 µl), consisting of 1 ml of 1M lithium acetate, 1 ml H_2_O and 8 ml PEG 3350 (50% w/v, freshly prepared and filtered), was added to the cells and was mixed. The cells were then subjected to heat shock at 42°C for 45 mins. Next, the cells were pelleted down, resuspended in 80µl of PBS solution, and were plated on plates containing Synthetic Defined medium without Histidine (SD-His). Transformed colonies obtained after 3-4 days were selected a second time on SD-His plates.

Genomic DNA from two colonies for each of the constructs were isolated. Cells from a colony were picked in 10µl sterile molecular grade water, and were incubated in a heat block at 70°C for 15min with 100µl of 200 µM lithium acetate and 1% (w/v) SDS. Genomic DNA was precipitated by adding 300µl of 100% Ethanol (Merck) and centrifuged at 15,000g for 5 min at room temperature. The supernatant was thrown away and pellet was dried. 50µl molecular grade water was added to the dried pellet, vortexed and centrifuged at 15,000g for 2 min. Thereafter, 40µl of supernatant was transferred to a fresh tube and 1 µl supernatant was taken for colony PCR to test the presence of the desired integrated fragment in the cells by PCR using Q5 DNA polymerase. PCR was performed using CHK primers (Tables S1) and the PCR program consisted of incubation at 98°C for 2 min, followed by 30 cycles of incubation at 98°C for 15s, at 60°C for 30s, at 72°C for 1 min, and a final incubation at 72°C for 5 min. The PCR products were confirmed by 1% agarose gel electrophoresis yielding product sizes of 792 bp, 786 bp and 690 bp for PDC1-GFP, ICL1-GFP and SOD1-GFP construct respectively.

### Gene expression quantification by flow cytometry

Expression levels of the tagged genes in all ancestral and evolved lines were quantified from two clones of each strain by flow cytometry. Verified GFP-tagged constructs and untagged BY4741 strain as the negative control were grown for 24 hours in test tubes containing 5ml YPD medium at 30°C with shaking at 200rpm. Cells were then diluted into fresh YPD medium (100 µl cells in 5 ml YPD) and were further grown for 4 hours to obtain cells in early log-phase of growth. 1ml of cells from 24h culture and 2ml of cells from 4h culture were taken for quantification. Cells were pelleted down, washed twice with PBS (8000 rpm, 5min) and resuspended in 1ml PBS. For 4h set, 1ml of resuspended cells and for 24 h set, 1ml of 4-fold diluted culture (250 µl culture + 750 µl PBS) were taken for flow cytometry experiments. In flow cytometry, 100,000 events for each sample were measured in both FITC and PE-Texas Red channels (Fig. S21).

The R package ‘flowCore’ was used to read fcs files and to convert them to text format. The data points with FSC-A value more than 2000 and SSC-A value more than 500 were considered as yeast cells. The points that showed lower values than these thresholds represented cell debris or noise and hence, were discarded. In addition, a filtering for cell aggregates were performed, since aggregates can generate stronger signal than single cells. To filter out cell aggregates, the ratio of FSC-A and FSC-H values were calculated, and data points showed two clusters. The data points with FSC-A/FSC-H value lower than 1.9 were considered as single cells and thus, were included in quantification of fluorescence signal (Fig. S22). The processed signal distribution from an evolved clone was compared with the results from a corresponding ancestral clone by Kolmogorov-Smirnov test. Thus, results for clones of lines N1, N2, and N3 were compared with that of clones of line A1, A3 and A3, respectively. Mode signal was estimated for unimodal distributions, whereas median signal was estimated for multi-modal distributions.

### Estimation of lipid content

Lipid content estimation in the ancestral and evolved lines was done using Nile red staining followed by quantification in flow cytometry. The staining technique was modified from Rostron and Lawrence, 2017. Nile Red stock solution at a concentration of 1mg/ml was prepared by dissolving Nile red powder in acetone. Ancestral and evolved lines were grown for ∼24 hours in 5ml YPD medium at 30°C with shaking at 200 rpm. Cells were then diluted 50-fold into fresh medium (100 µl cells in 5ml YPD) and were grown for 6 hours at 30°C, 200 rpm to reach the early-to mid-log phase of growth. 100 μl cells from saturated cultures (after 24h of growth) and 1ml cells from cultures at 6h were taken, washed with PBS and finally resuspended in 1 ml PBS. For staining, cells were incubated with 1 μl Nile Red (stock 1 mg/ml) at a final concentration of 1 μg/ml at room temperature in the dark for 5 mins. Cells were then washed twice with PBS, and resuspended in 1 ml PBS for flow cytometry (Fig. S23). Confocal microscopy (FLUOVIEW FV3000 laser scanning microscope, Olympus) images were also captured for visualization of the lipid droplets at 100X magnification using Alexa Fluor 488 and yellow-red filters. Flow cytometry data were analyzed following the same method described before. The processed signal distribution from an evolved line was compared with the results from the ancestral line by Kolmogorov-Smirnov test. Thus, the results from N1, N2 and N3 lines were compared with that of A1, A2 and A3 lines respectively.

### Selection experiment in new environments

All three replicates of the evolved lines along with the replicates of ancestral lines were selected in new environmental conditions. The selection conditions included minimal growth medium (SCD), SCD + 1M NaCl, SCD + 4mM H_2_O_2_, SCD + 0.08 µg/ml caspofungin, and SCD + 3 µg/ml flucytosine. Selection was done in 96-deepwell plates where the total culture volume in each well was 1.2ml, and the plates were sealed with Aeraseal (Sigma) to allow air exchange. Cells from each line were grown overnight in SCD medium and ∼10^5^ cells were put in each selection condition and were allowed to grow for 48 hours at 30°C with shaking, thereby completing one round of selection. After 48 hours, ∼10^5^ cells were put in fresh medium with identical selection condition, and were allowed to grow again for 48 hours. This process was repeated for 10 rounds. Cell densities were estimated at the end of each round through measurement of optical density.

### Genome sequencing

Genomic DNA of all ancestral (A) and evolved (N) lines were isolated using Yeast DNA Extraction kit (Thermo Scientific) following manufacturer’s protocol. Briefly, overnight grown yeast cells were treated with lysing and protein removal agents (provided in the kit). The genomic DNA was then precipitated with isopropyl alcohol, the pellet washed with 70% ethanol and finally resuspended in TE buffer (10 mM Tris), yielding about 35-70 ng/µl DNA for each line. Sequencing was done using the Illumina Novaseq 6000 platform to obtain ∼5 million paired-end 150 bp (2 × 150bp) reads for each sample.

### Variant identification

Paired-end read data for each line was obtained as fastq files. Quality control of the data was done using the ‘FastQC’ tool (https://www.bioinformatics.babraham.ac.uk/projects/fastqc/). Data showed adapter contamination, which were subsequently trimmed by *bbduk* function from ‘bbmap’ package (BBMap - Bushnell B. - sourceforge.net/projects/bbmap/). The reads were then mapped to the reference *S. cerevisiae* genome (version R64.2.1) using the package *bowtie2* (Langmead and Salzberg 2012) with *--local* option. The output file in *sam* format was further processed using custom codes as follows. Only concordantly uniquely mapped reads were considered, and were locally realigned against the mapped region to ensure proper alignment. After realignment, mutations including SNPs and indels were noted down including the quality of the mutated base. In the next step, paired-end reads were merged to generate a consolidated list of mutations. The reads were first checked for any overlap in between. If a mutation appeared in only one of the reads in the overlapping region, it was considered a sequencing error and hence was discarded. Mutations appearing in non-overlapping regions were carried forward for further analysis. The merged read data was also used for calculating sequencing coverage for each position in the genome in a sample. For an overlapping region between two reads, sequencing coverage for every base increased by a value of one, similar to that of non-overlapping regions.

In the next step, sequencing coverage for each position of the genome was computed. The mutation data obtained from the merging step of paired-end reads was used for a definite identification of mutation and for calculating their frequency in populations. A mutation was considered a true mutation if it was present in at least 10 reads covering that position, and if it fulfilled at least one of the following conditions. The mutation had an average quality score of at least 30, or had quality score of at least 20 and supported by both read pairs, or was a deletion mutant that was present in both read pairs. Finally, mutations were compared among ancestral and evolved lines, and the mutations were annotated according to their location in a gene or promoter region, and the type of coding mutation, such as synonymous, nonsynonymous and nonsense mutations.

To test whether a mutation observed in the lines selected in antifungals, starting from either N lines or A lines, was also present in any read of the genomic data of the corresponding N line or A line from which selection experiments were initiated, a search was performed for the same mutations in the reads without any consideration for quality score of the mutated base. Only the number of non-duplicate reads containing the mutation were counted. The enrichment value for each mutation was calculated as follows.

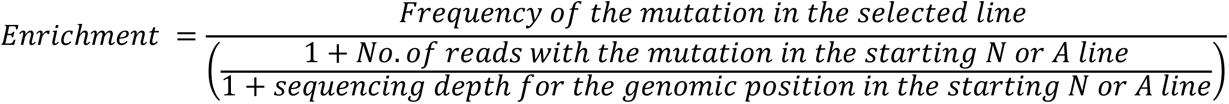

### Copy number variation (CNV) analysis

Sequencing coverage for every position in the genome in all lines were first quantified, and were normalized by median-of-ratios method. First, for every position, the geometric mean of the coverage across all samples was calculated. In the next step, ratio of the sequencing coverage and the geometric mean was calculated. Finally, the median value of the ratios for all genomic positions in a sample was considered as the normalization factor. For each position in a sample, the normalization factor was used to normalize coverage data. Subsequently, normalized log_2_ ratio for each genomic position was calculated from a ratio between normalized coverage in an evolved sample and normalized cover in an ancestral sample. The ratio was finally reported as 1 + Normalized log_2_ coverage ratio. A value of 1 suggested no change in copy number, a value of 2 or more suggested segmental amplifications, whereas a value of 0 suggested deletion. Regions of at least 1 kb length and with 1+normalized log2 coverage ratio scores above 1.5 or below 0.2 were identified as amplifications or deletions respectively.

## Acknowledgments

The authors are thankful to the members of the lab of Dr. Gayatri Mukherjee, School of Medical Science and Technology, IIT Kharagpur for help with flow cytometry. The authors also acknowledge support from the University of Exeter Sequencing facility. This work utilized equipment funded by the DST-FIST, Govt. of India, and the Wellcome Trust (Multi-User Equipment Grant award number 218247/Z/19/Z).

The authors are grateful to Dr. Dragan Stajic for a critical reading of the first draft of the manuscript.

## Author contributions

Conceptualization: RD

Methodology: SR

Investigation: SR, RL

Visualization: SR, RD

Funding acquisition: IG, RD

Project administration: IG, RD

Supervision: IG, RD

Writing – original draft: SR, RL, IG, RD

Writing – review & editing: SR, RL, IG, RD

