## Supplementary figures and tables for "Exaptation and de novo mutations transcend cryptic variations as drivers of adaptation in yeast"

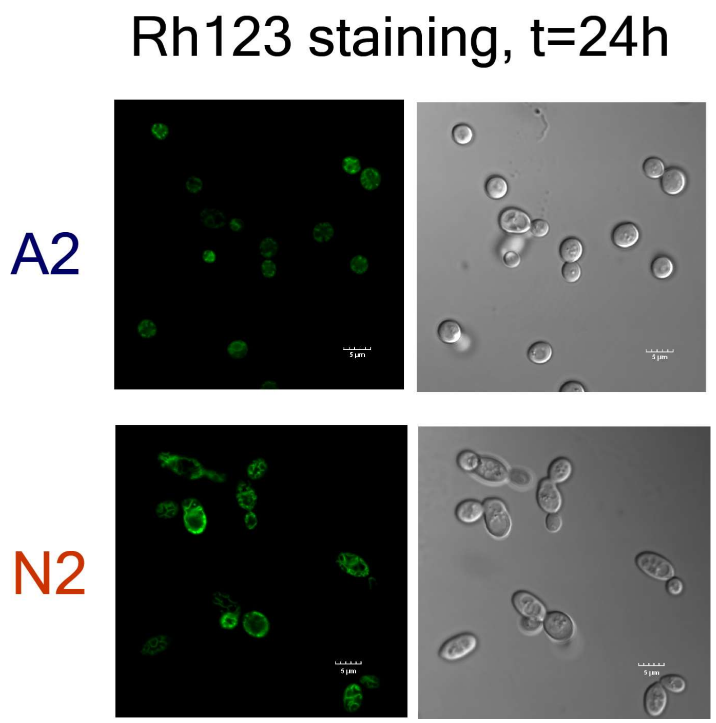


**Figure S1** - Microscopic images showing mitochondrial network in A2 and N2 lines stained with Rhodamine 123 at 24 hours of growth.

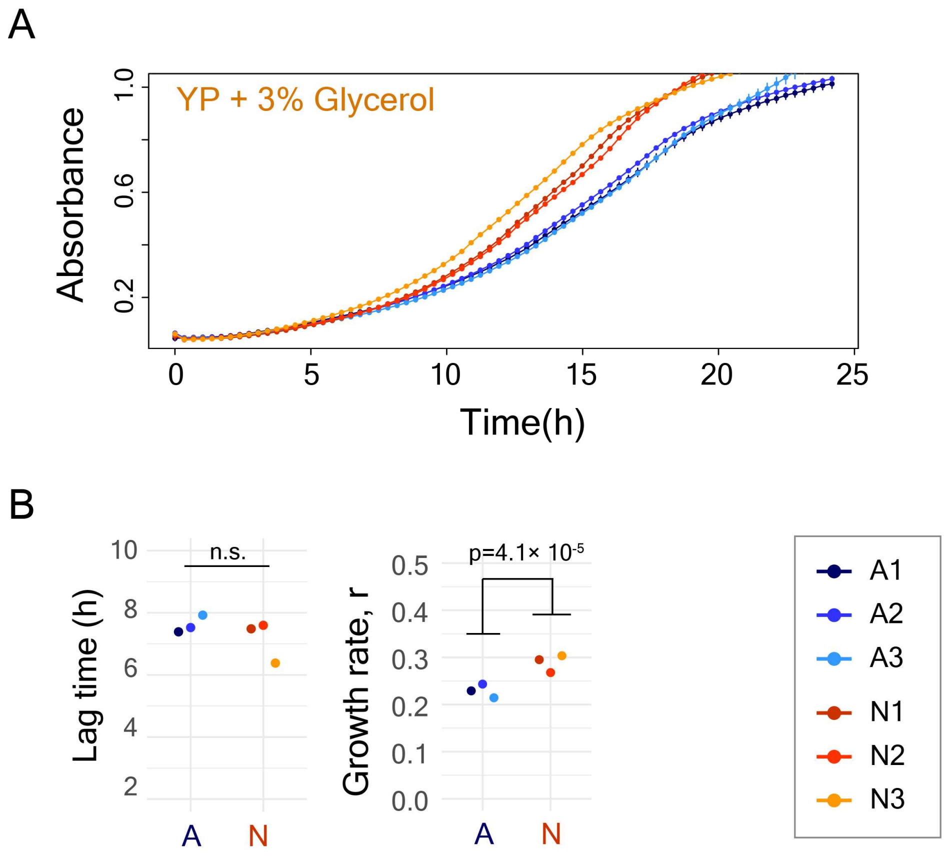


**Figure S2** – **(A)** Growth curve of ancestral and evolved lines in YP medium supplemented with 3% glycerol. **(B)** Estimates of lag-time and growth rate from growth curves in YP+3% glycerol.

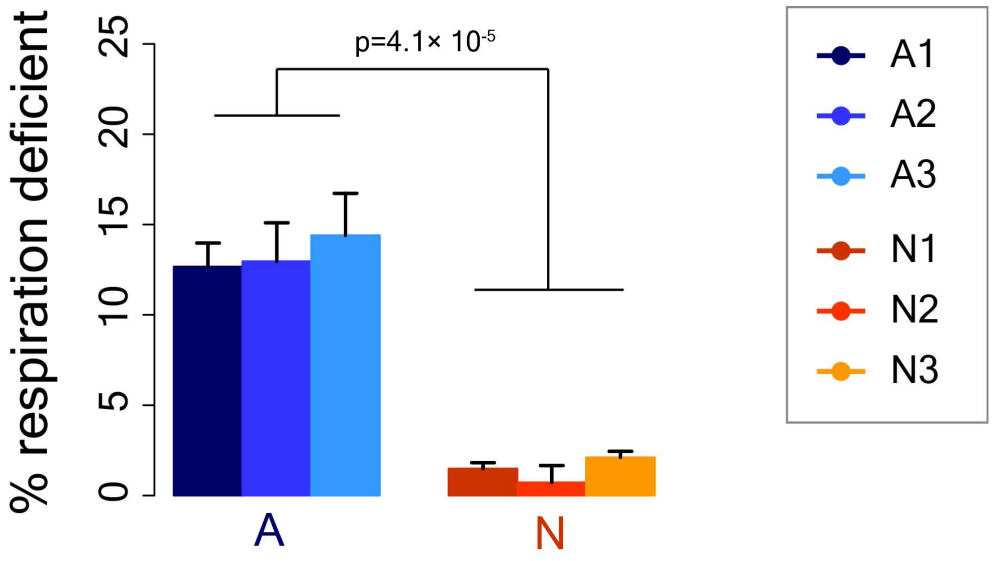


**Figure S3** - Percentage of respiration deficient cells in the ancestral and the evolved lines.

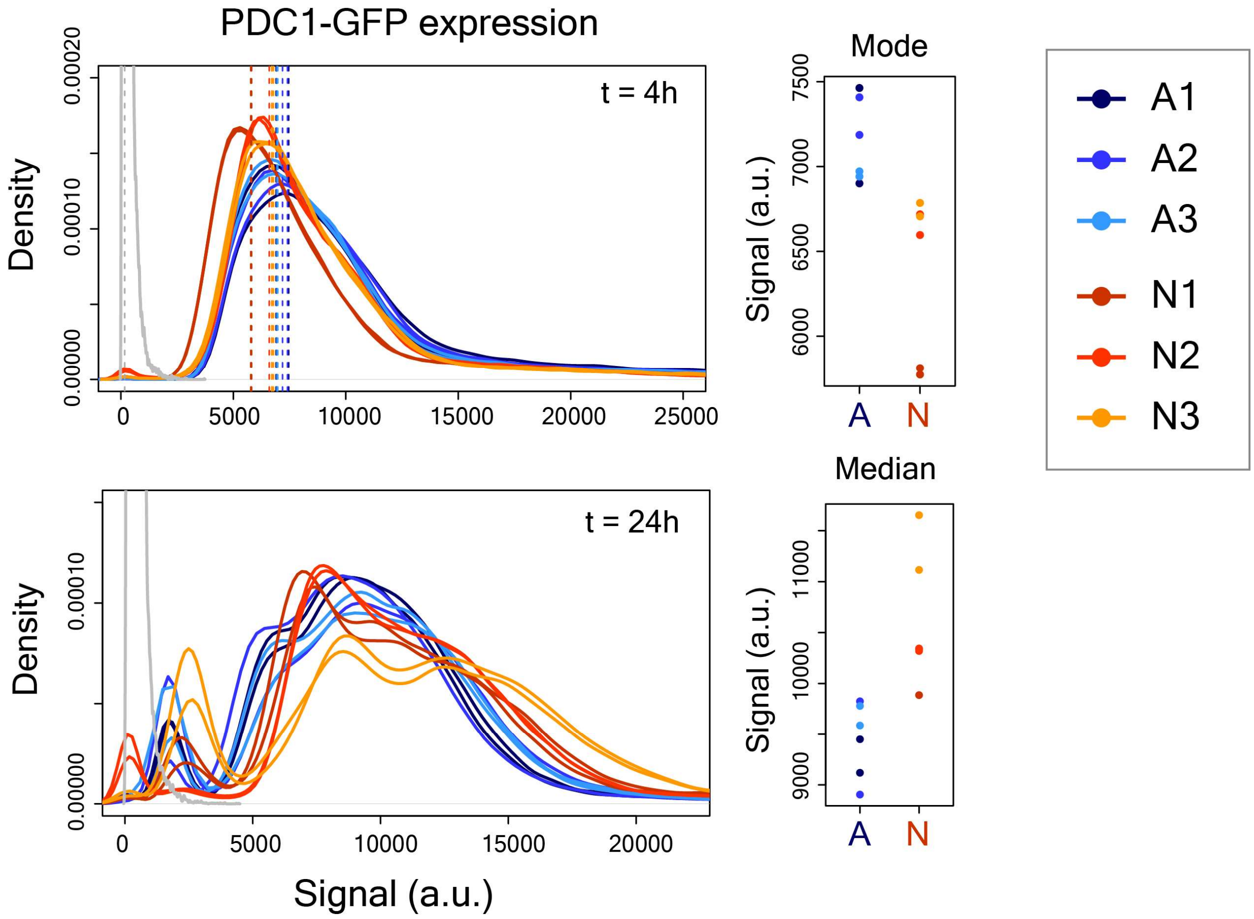


**Figure S4** - Expression of PDC1-GFP fusion protein measured by flow cytometry at 4 hours and 24 hours of growth (Kolmogorov-Smirnov test for comparison between clones of corresponding evolved and ancestral lines, $p<10^{-10}$ at t=4h and at t=24h). Mode signal was estimated for unimodal distributions, whereas median signal was estimated for multi-modal distributions.

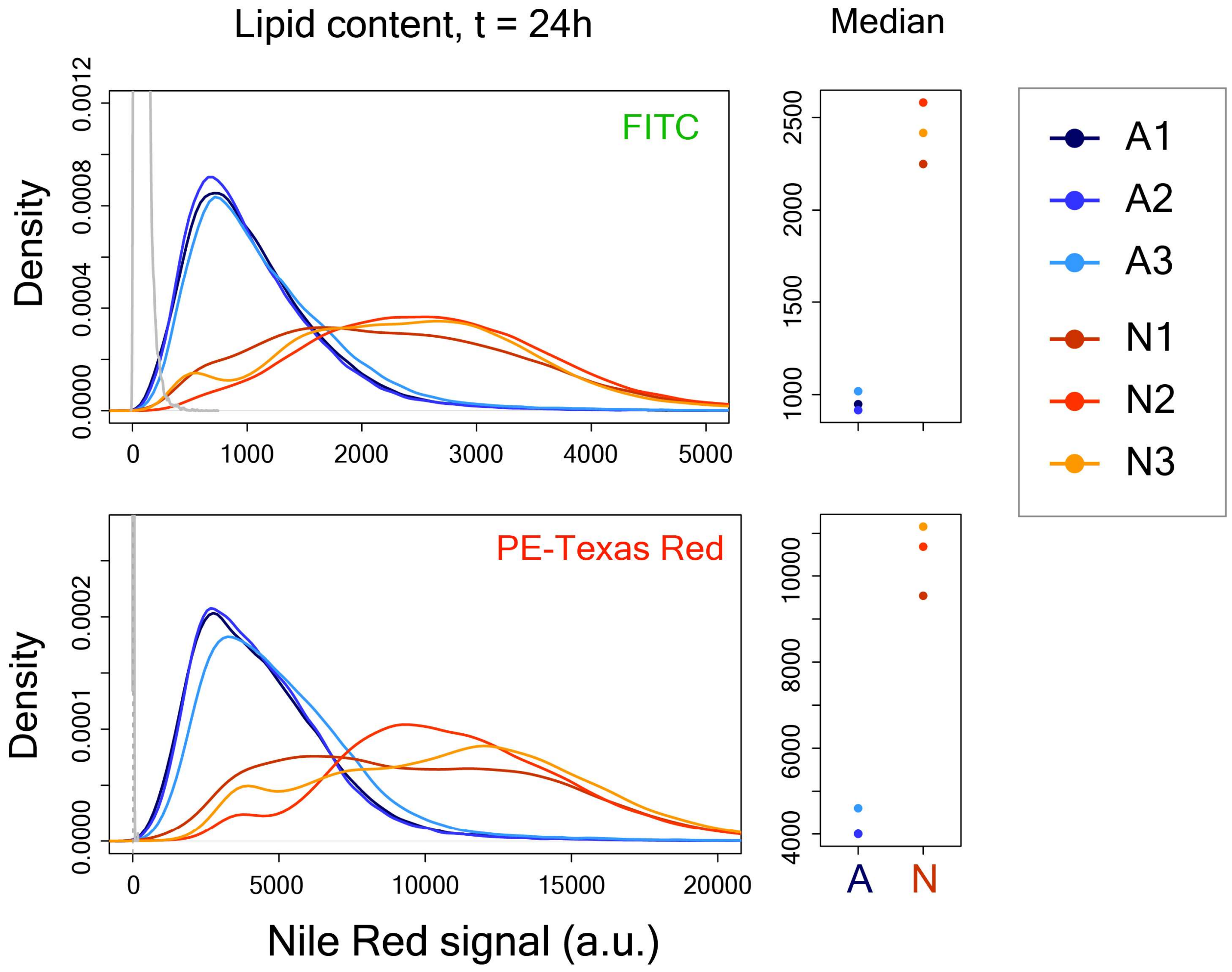


**Figure S5** - Lipid content per cell in ancestral and evolved lines after 24 hours of growth quantified through flow cytometry using the Nile red stain.

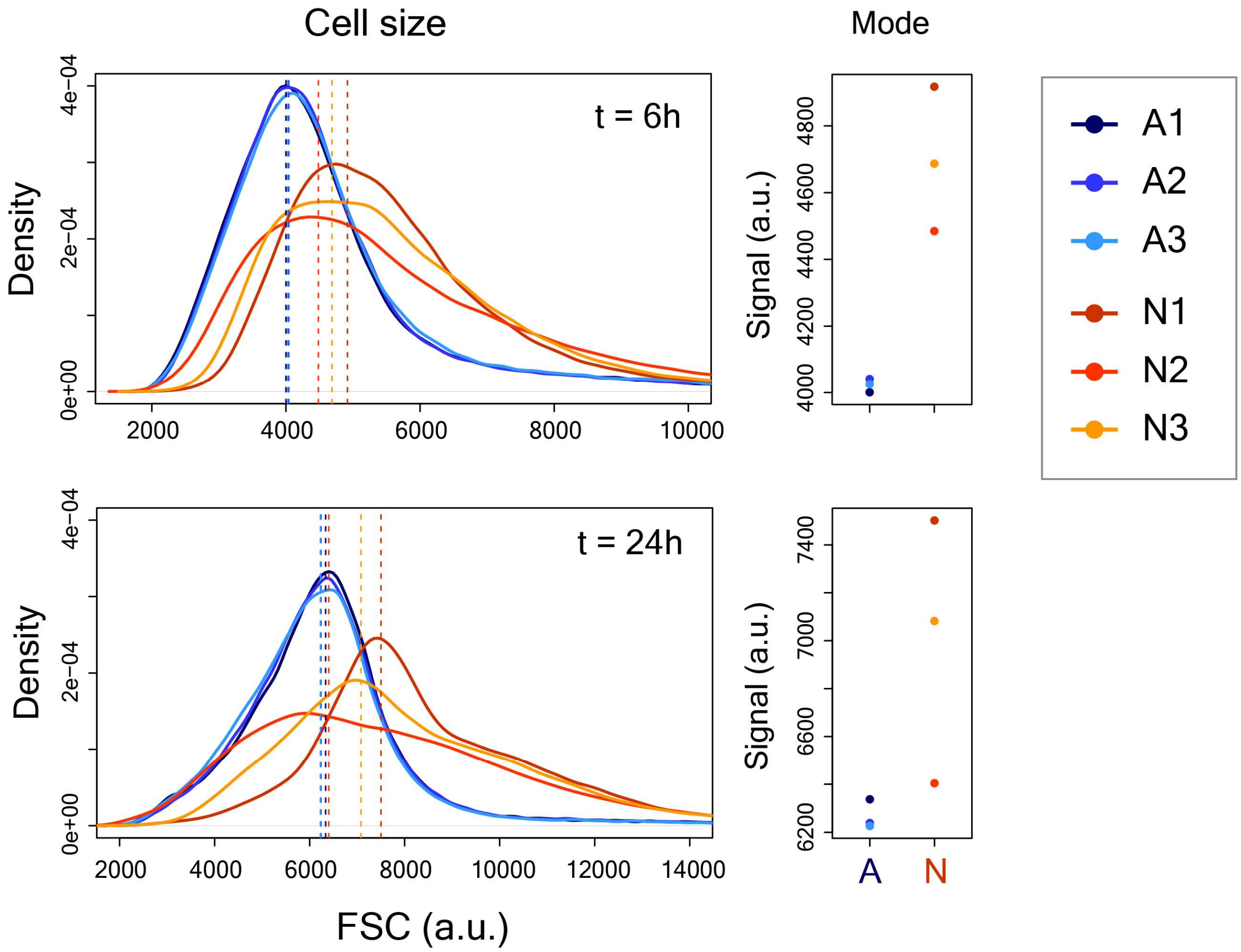


**Figure S6** - Increased cell size in evolved lines as observed in flow cytometry at 6 hours and 24 hours of growth (Kolmogorov-Smirnov test for comparison between corresponding evolved and ancestral lines, $p<10^{-10}$ for all comparisons).

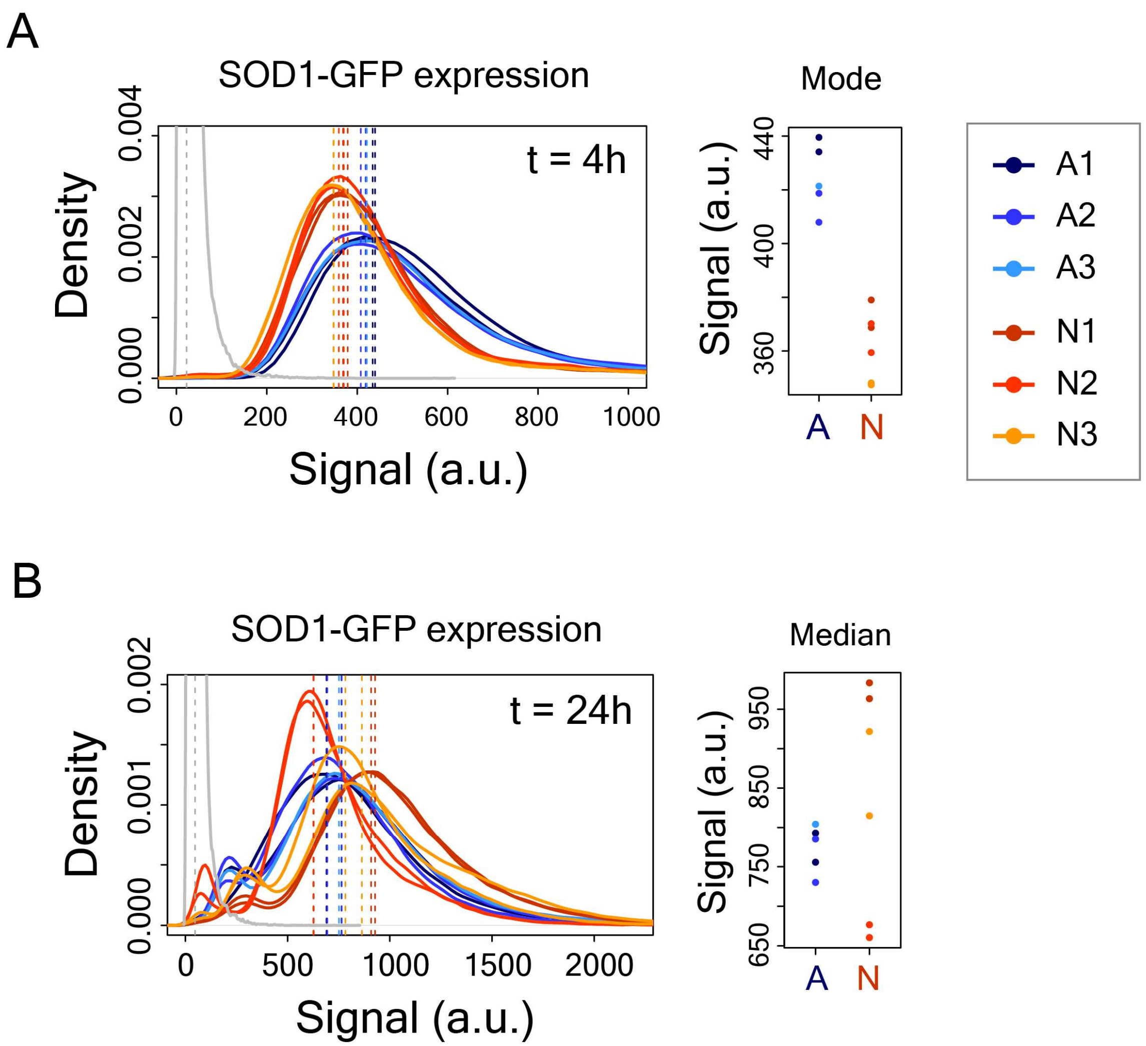


**Figure S7** - **(A)** SOD1-GFP expression quantified by flow cytometry at 4 hours of growth **(B)** SOD1-GFP expression quantified by flow cytometry at 24 hours of growth. The distributions in grey shows data from negative controls without any GFP tag or without staining. ($p<10^{-10}$ for each comparison, Kolmogorov-Smirnov test between signal distributions of clones of corresponding evolved and ancestral lines). Mode signal was estimated for unimodal distributions, whereas median signal was estimated for multi-modal distributions. The dotted lines show mode (A) or median (B) values.

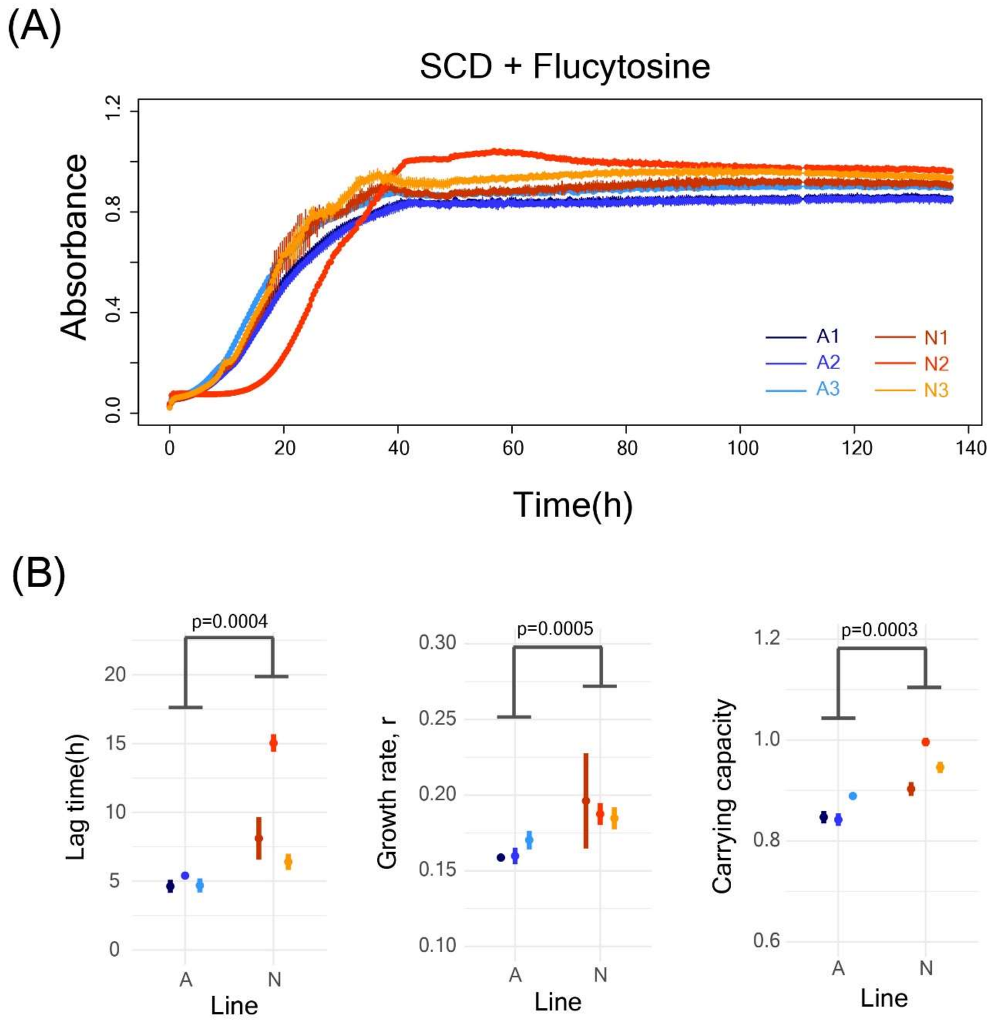


**Figure S8 - (A)** Growth curves of the ancestral and evolved lines in SCD medium supplemented with 3 µg/ml flucytosine. **(B)** Growth parameters estimated from the growth curves in (A).

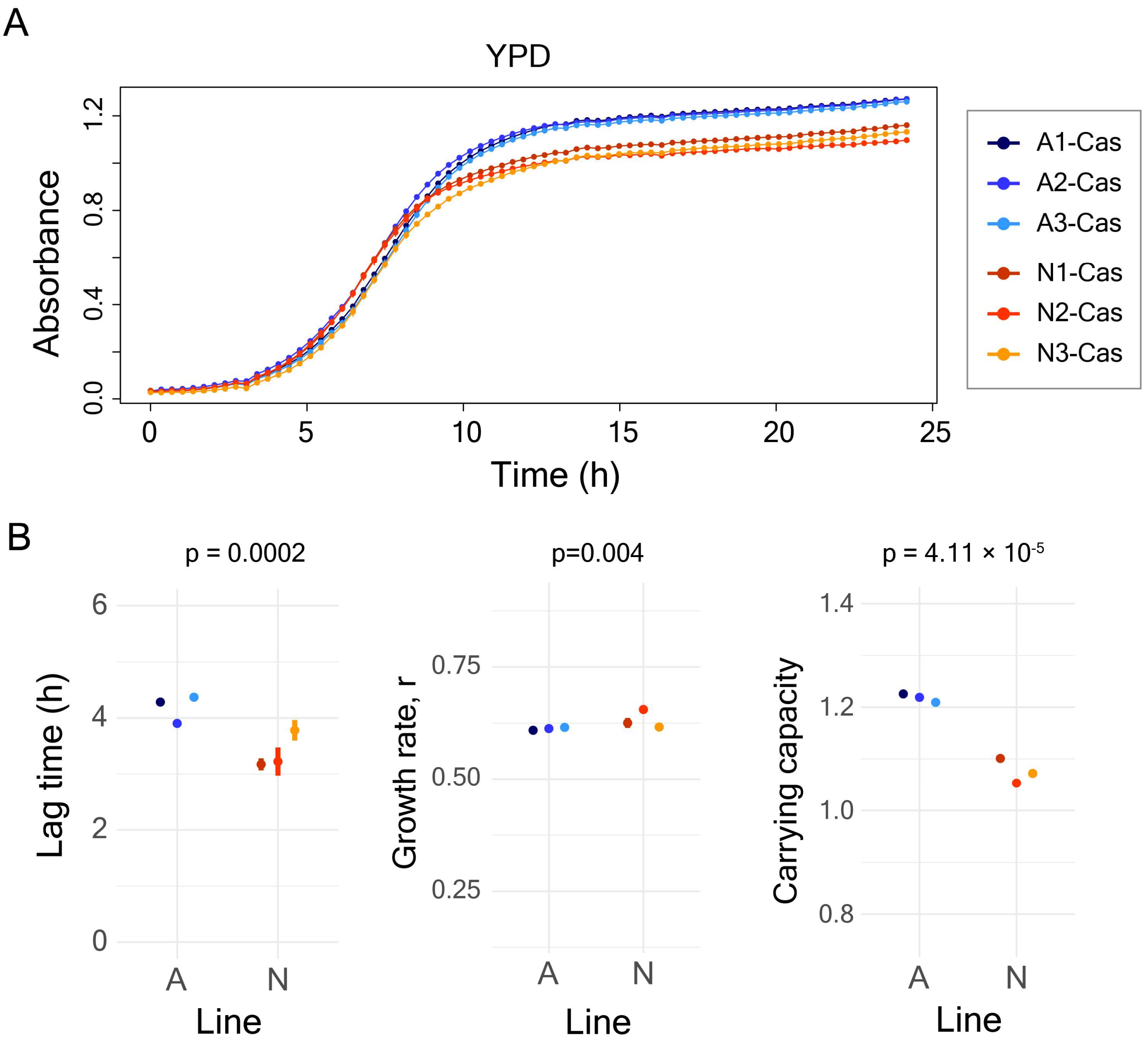


**Figure S9 - (A)** Growth curves in YPD medium of the A and N lines selected in SCD + 0.08 µg/ml caspofungin for 10 rounds **(B)** Growth parameters estimated from the growth curves in (A).

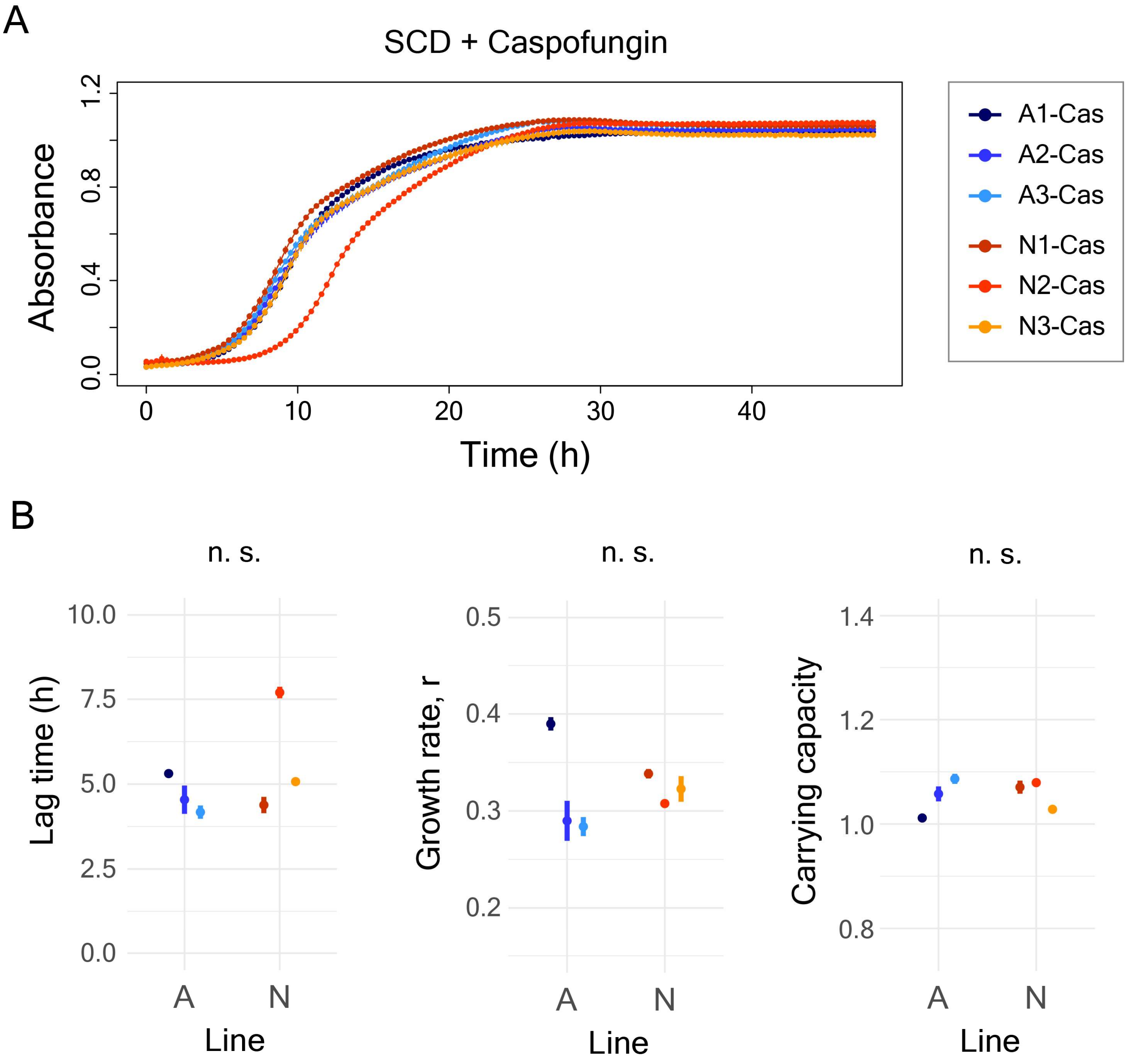


**Figure S10 - (A)** Growth curves in SCD medium with 0.08 µg/ml caspofungin of the A and N lines selected for 10 rounds in SCD + 0.08 µg/ml caspofungin **(B)** Growth parameters estimated from the growth curves in (A).

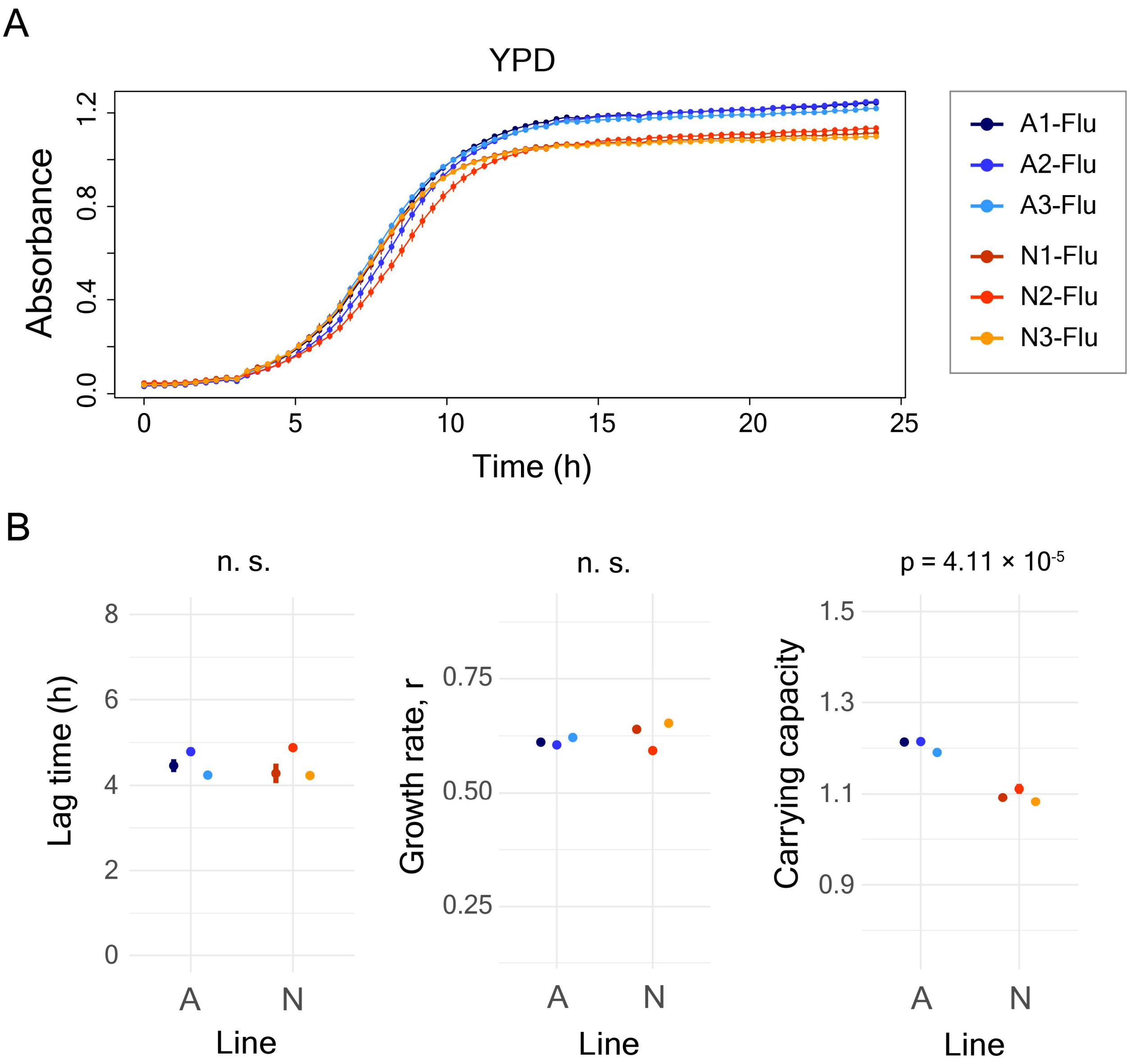


**Figure S11 - (A)** Growth curves in YPD medium of the A and N lines selected in SCD + 3 µg/ml flucytosine for 10 rounds **(B)** Growth parameters estimated from the growth curves in (A).

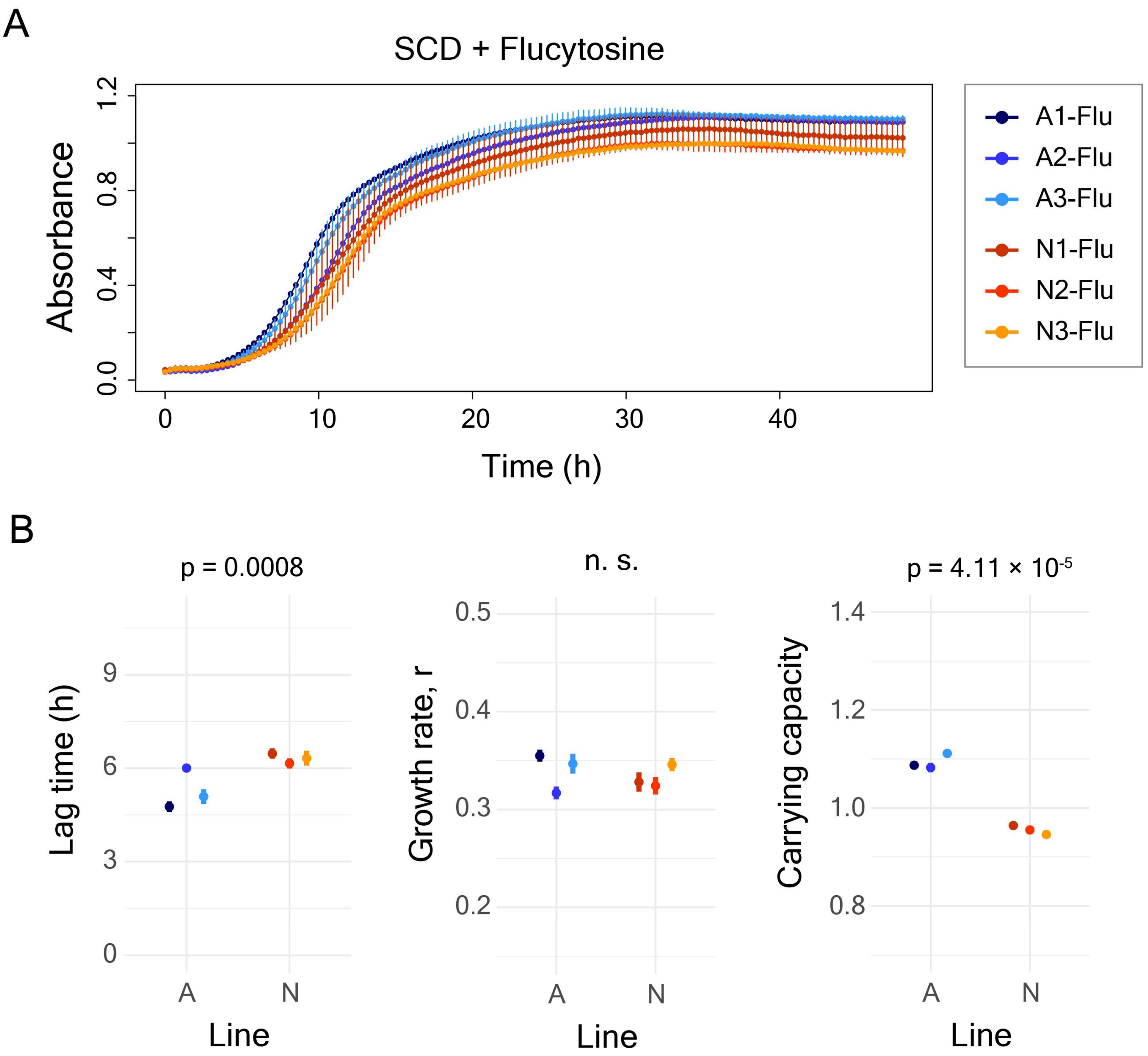


**Figure S12 - (A)** Growth curves in SCD medium with 3 μg/ml flucytosine of the A and N lines selected in SCD + 3 μg/ml flucytosine for 10 rounds **(B)** Growth parameters estimated from the growth curves in (A).

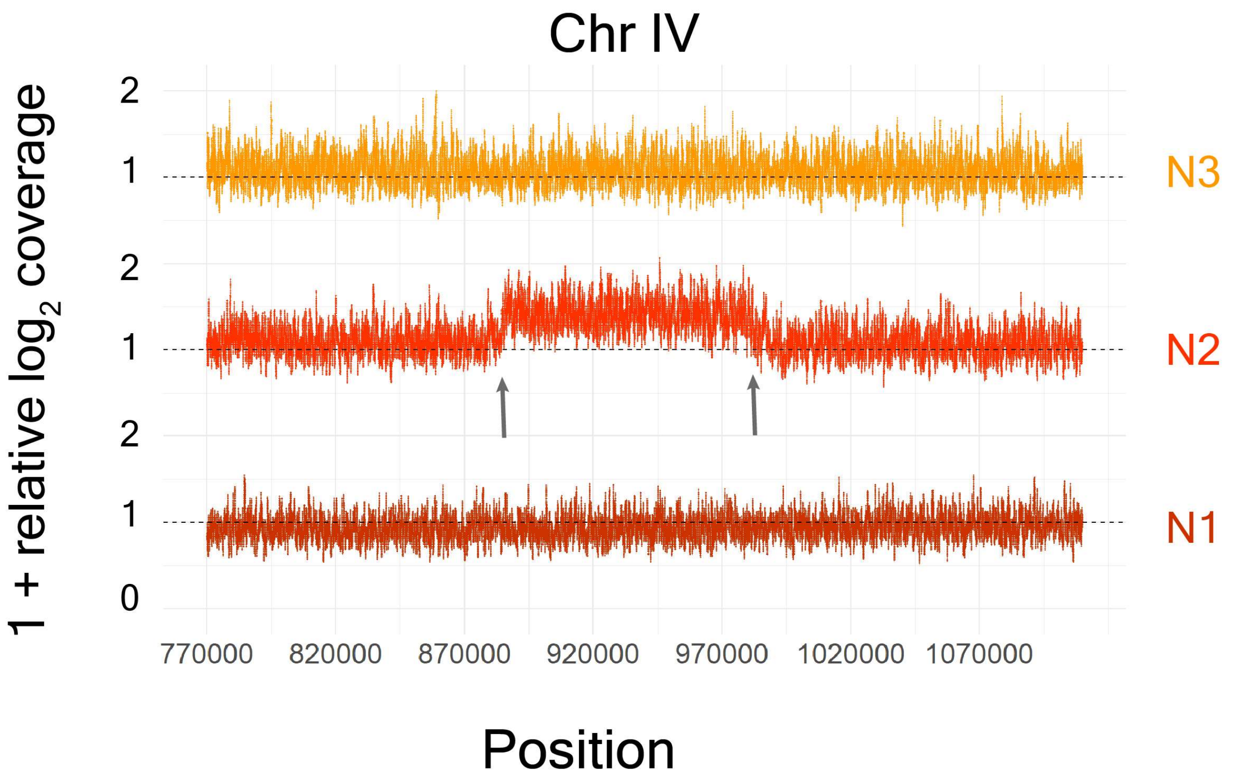


**Figure S13** – CNV analysis of the N lines compared to the A lines revealed a segmental amplification in ChrIV of the N2 line. The x-axis shows positions in ChrIV and the y-axis shows (1 + relative $\log_2$ coverage) values**.**

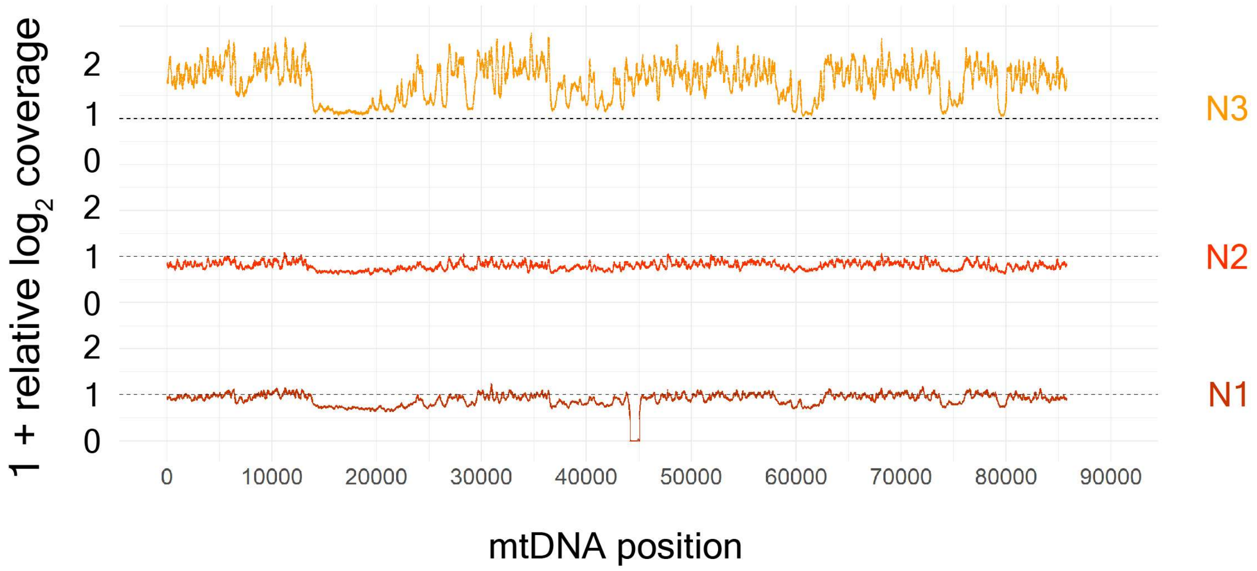


**Figure S14** – CNV analysis reveals an increase in mtDNA copy number in the N3 line. The x-axis shows positions in mtDNA and the y-axis shows (1 + relative $\log_2$ coverage) values**.**

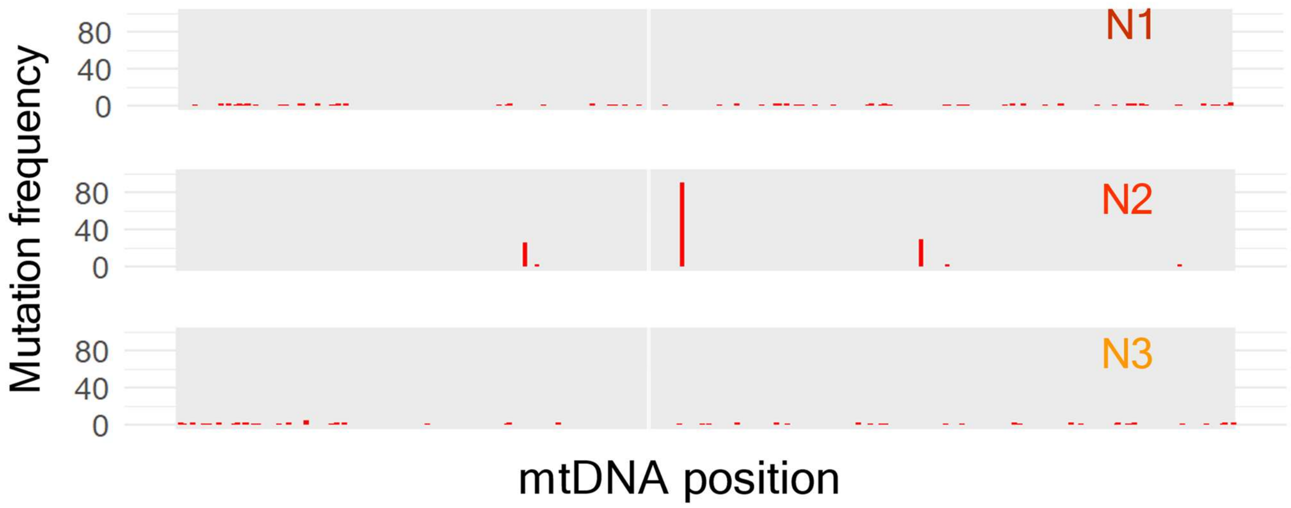


**Figure S15** – Mutations observed in mtDNA along with their frequencies in the N lines

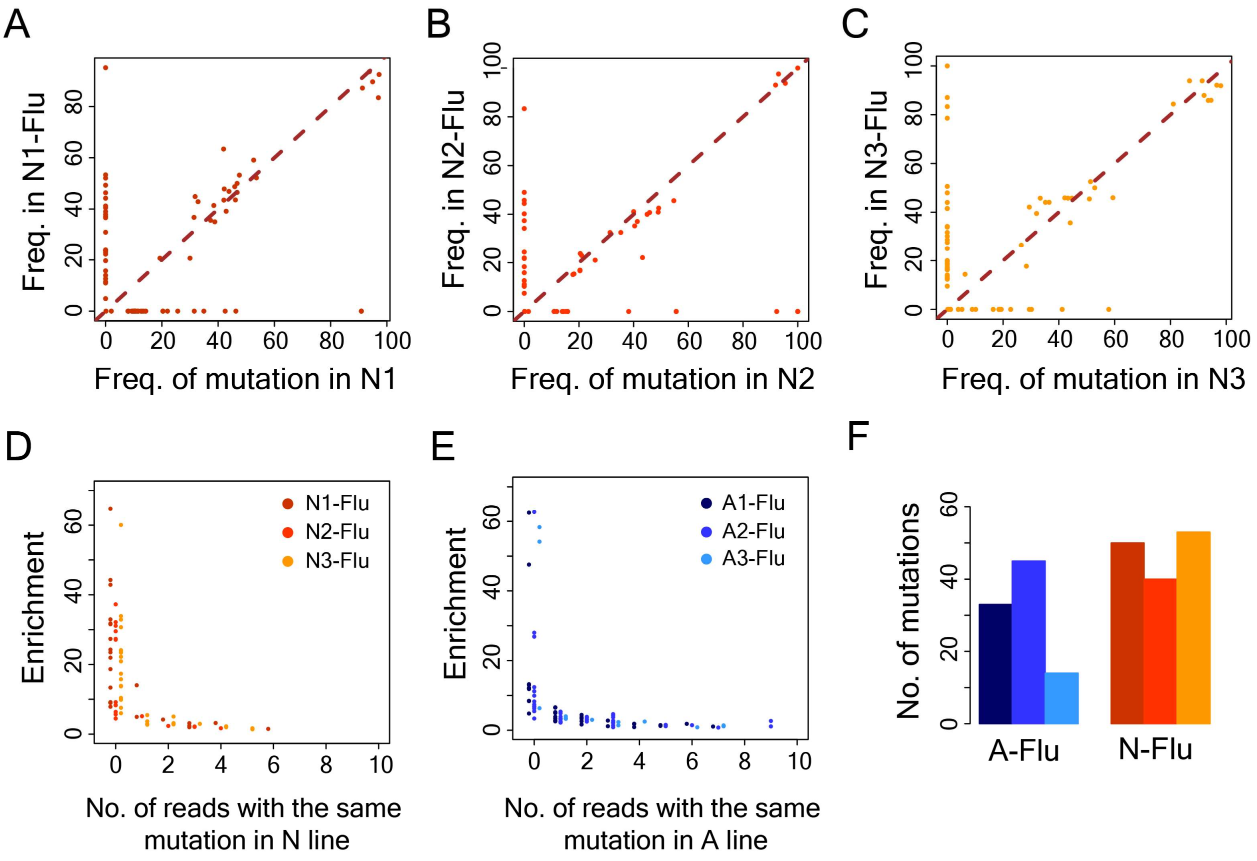


**Figure S16 - Adaptation to flucytosine is driven by de novo mutations. (A)** Frequency of mutations in the N1 line before (N1) and after (N1-Flu) flucytosine selection **(B)** Frequency of mutations in the N2 line before (N2) and after (N2-Flu) flucytosine selection **(C)** Frequency of mutations in the N1 line before (N3) and after (N3-Flu) flucytosine selection **(D)** The y-axis shows enrichment of mutations in the flucytosine-selected N lines and the x-axis shows the number of nonduplicate reads in the genomic data of the corresponding N line in which the mutation was detected **(E)** The y-axis shows enrichment of mutations in the flucytosine-selected A lines and the x-axis shows the number of nonduplicate reads in the genomic data of the corresponding A line in which the mutation was detected **(F)** Distribution of number of mutations in flucytosine-selected A and N lines (denoted by A-Flu and N-Flu, respectively).

BD FACSDiva 8.0.2

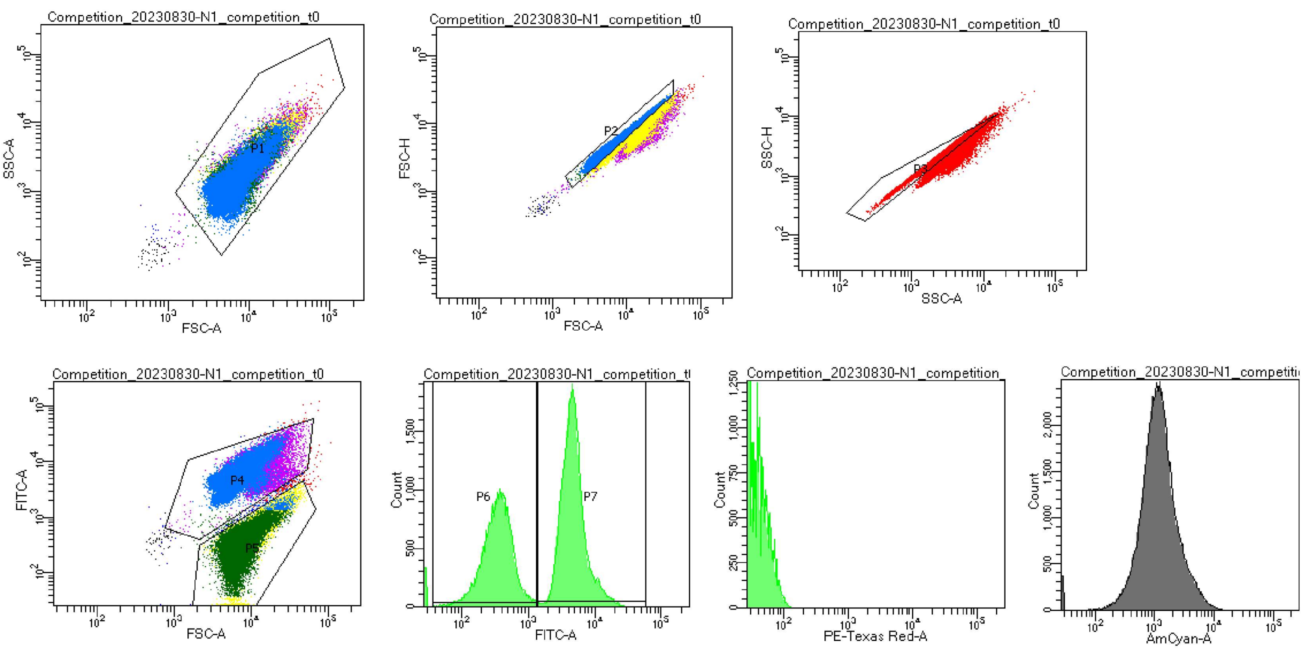


Tube Name: N1_competition_t0
GUID: 48d0b905-0e45-485d-982d-b2ed0e98fa0d

| Population | #Events | %Parent | FSC-A Mean | SSC-A Mean | FITC-A Mean | FITC-A Median | FITC-A SD | FITC-A %CV | PE-Texas ... Mean | PE-Texas ... Median | AmCyan-A Mean | AmCyan-A Median |
|---|---|---|---|---|---|---|---|---|---|---|---|---|
| All Events | 100,000 | #### | 8,215 | 1,950 | 3,680 | 3,161 | 4,253 | 115.6 | 9 | 8 | 1,442 | 1,078 |
| P1 | 99,883 | 99.9 | 8,222 | 1,952 | 3,683 | 3,165 | 4,254 | 115.5 | 9 | 8 | 1,441 | 1,077 |
| P2 | 77,465 | 77.5 | 6,835 | 1,523 | 3,086 | 2,905 | 3,287 | 106.5 | 7 | 6 | 1,124 | 946 |
| P3 | 87,449 | 87.4 | 7,018 | 1,637 | 3,186 | 2,853 | 3,523 | 110.6 | 8 | 7 | 1,203 | 986 |
| P4 | 58,097 | 58.1 | 7,893 | 1,771 | 5,974 | 4,762 | 4,057 | 67.9 | 8 | 7 | 1,575 | 1,177 |
| P5 | 41,494 | 41.5 | 8,532 | 2,146 | 434 | 353 | 349 | 80.3 | 11 | 9 | 1,205 | 950 |
| P6 | 32,637 | 42.1 | 7,530 | 1,793 | 362 | 328 | 196 | 54.2 | 9 | 8 | 985 | 874 |
| P7 | 44,350 | 57.3 | 6,315 | 1,323 | 5,122 | 4,372 | 3,024 | 59.0 | 6 | 5 | 1,229 | 1,003 |

**Figure S17** – A snapshot of flow cytometry experiment at t=0 for competition assay

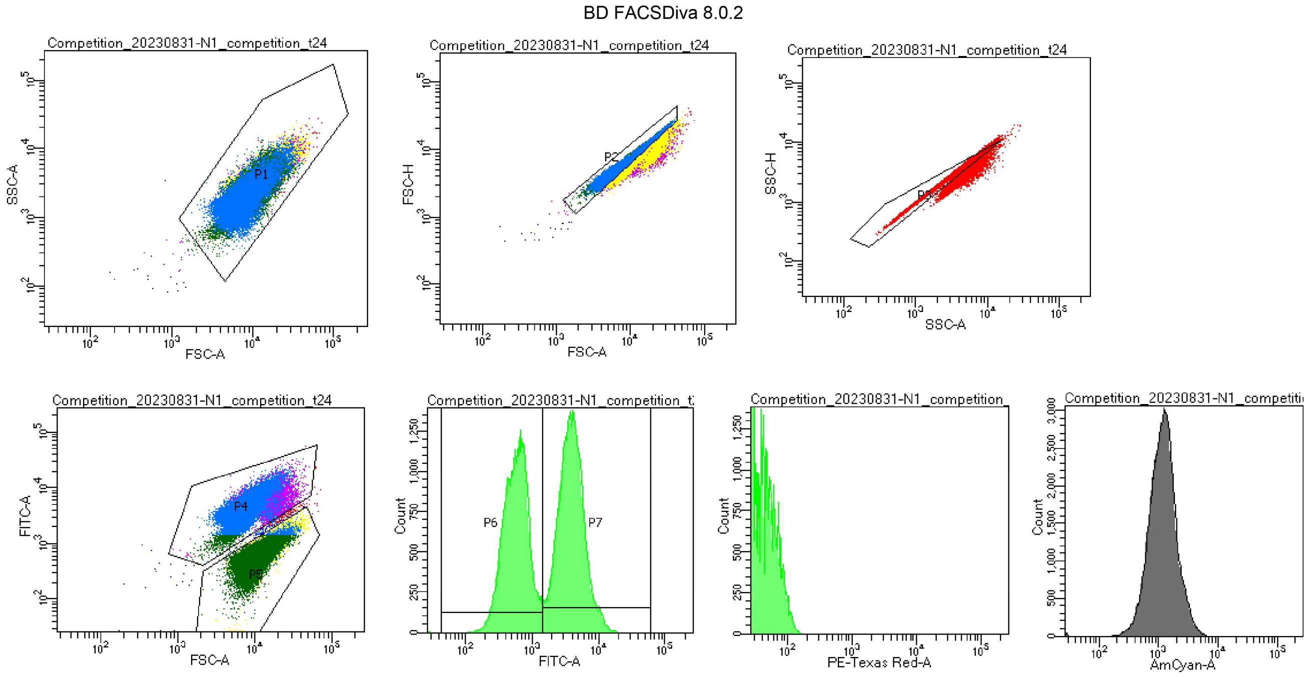


| Tube Name: | | | N1_competition_t24 | | | | | | | | |
|---|---|---|---|---|---|---|---|---|---|---|---|
| GUID: | | | d1fbe22e-d94f-4e01-a6cb-e13a56156bf2 | | | | | | | | |
| Population | #Events | %Parent | FSC-A Mean | SSC-A Mean | FITC-A Mean | FITC-A Median | FITC-A SD | FITC-A %CV | PE-Texas ... Mean | PE-Texas ... Median | AmCyan-A Mean | AmCyan-A Median |
| All Events | 100,000 | #### | 10,079 | 2,686 | 2,576 | 1,687 | 2,591 | 100.6 | 18 | 16 | 1,256 | 1,103 |
| P1 | 99,950 | 100.0 | 10,083 | 2,688 | 2,577 | 1,690 | 2,591 | 100.5 | 18 | 16 | 1,256 | 1,103 |
| P2 | 76,049 | 76.0 | 9,252 | 2,395 | 2,459 | 1,886 | 2,353 | 95.7 | 16 | 14 | 1,132 | 1,019 |
| P3 | 94,083 | 94.1 | 9,419 | 2,499 | 2,434 | 1,388 | 2,419 | 99.4 | 17 | 15 | 1,170 | 1,065 |
| P4 | 50,638 | 50.6 | 8,578 | 2,080 | 4,474 | 3,969 | 2,396 | 53.6 | 12 | 10 | 1,190 | 1,020 |
| P5 | 48,992 | 49.0 | 11,550 | 3,289 | 613 | 553 | 309 | 50.4 | 24 | 21 | 1,312 | 1,187 |
| P6 | 35,865 | 47.2 | 10,879 | 3,004 | 579 | 544 | 246 | 42.4 | 22 | 20 | 1,196 | 1,136 |
| P7 | 40,163 | 52.8 | 7,801 | 1,851 | 4,139 | 3,688 | 2,108 | 50.9 | 11 | 9 | 1,075 | 936 |

**Figure S18 –** A snapshot of flow cytometry experiment at t=24h for competition assay

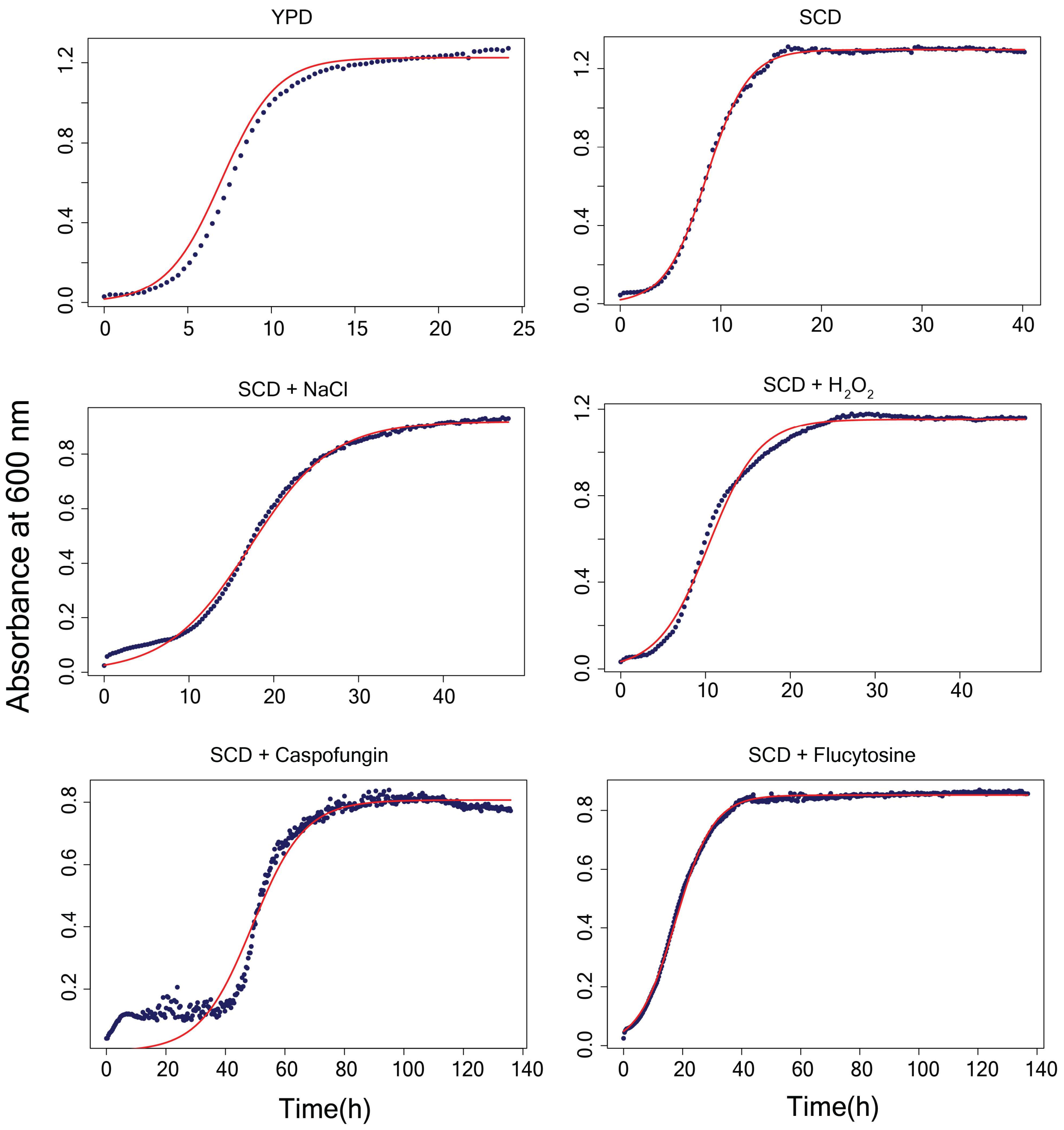


**Figure S19 – Fitting of growth curves using logistic growth equation to estimate growth parameters (growth rate, initial cell density and carrying capacity).** The blue points show actual observations from the growth curve of the ancestral line A1 in SCD medium, whereas the red line shows a line fitted with the estimated growth parameters from that growth curve.

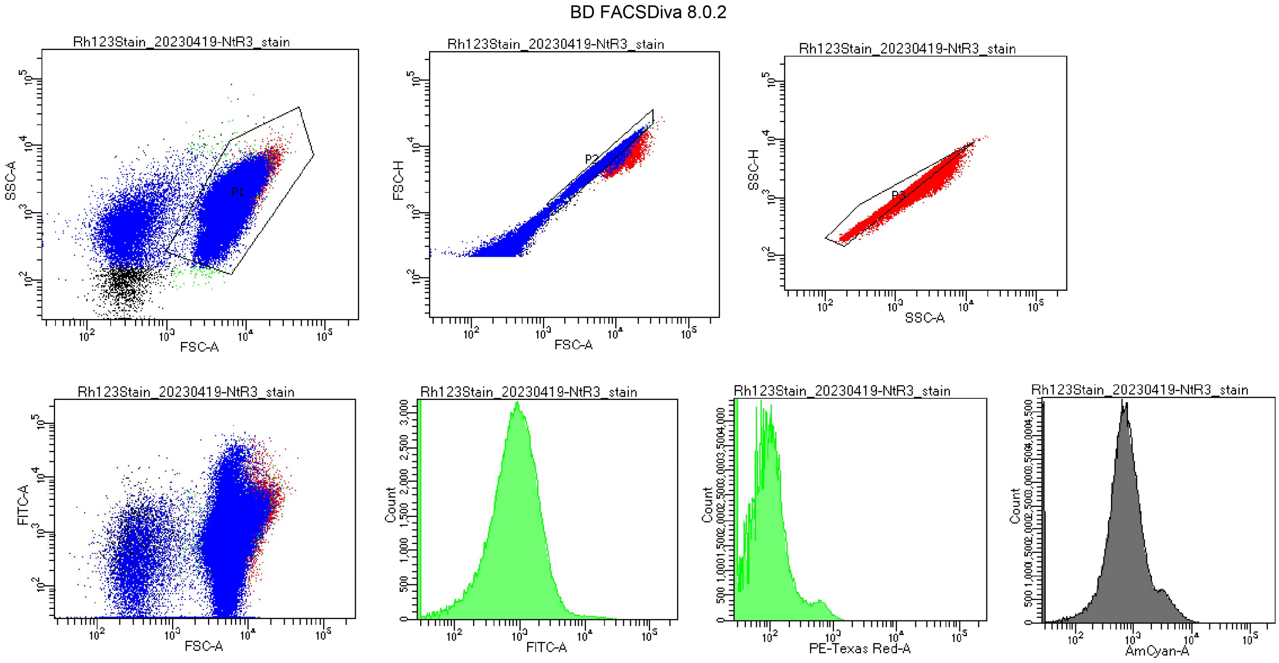


Tube Name: NtR3_stain
GUID: c50eedc2-49df-4854-baed-d0dc8afa15e1

| Population | #Events | %Parent | FSC-A Mean | SSC-A Mean | FITC-A Mean | FITC-A Median | FITC-A SD | FITC-A %CV | PE-Texas ... Mean | PE-Texas ... Median | AmCyan-A Mean | AmCyan-A Median |
|---|---|---|---|---|---|---|---|---|---|---|---|---|
| All Events | 200,000 | #### | 5,916 | 1,536 | 1,190 | 778 | 2,183 | 183.5 | 118 | 79 | 1,012 | 670 |
| P1 | 188,260 | 94.1 | 6,255 | 1,579 | 1,237 | 813 | 2,216 | 179.1 | 123 | 83 | 1,061 | 698 |
| P2 | 178,818 | 89.4 | 5,902 | 1,490 | 1,165 | 777 | 2,130 | 182.7 | 116 | 80 | 999 | 673 |
| P3 | 186,203 | 93.1 | 5,628 | 1,450 | 1,122 | 745 | 2,101 | 187.2 | 111 | 77 | 957 | 649 |

**Figure S20 –** A snapshot of flow cytometry experiment after Rh123 staining

BD FACSDiva 8.0.2

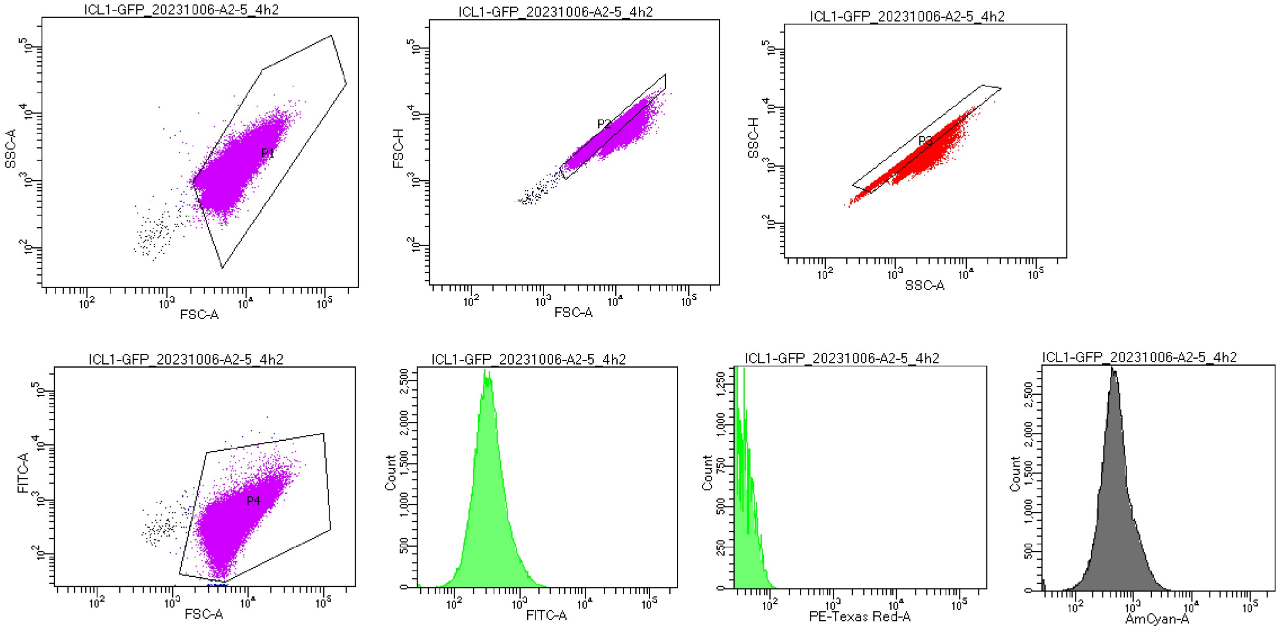


Tube Name: A2-5_4h2
GUID: 6e9cad75-1250-423a-b6e9-84cc8a611425

| Population | #Events | %Parent | FSC-A Mean | SSC-A Mean | FITC-A Mean | FITC-A Median | FITC-A SD | FITC-A %CV | PE-Texas ... Mean | PE-Texas ... Median | AmCyan-A Mean | AmCyan-A Median |
|---|---|---|---|---|---|---|---|---|---|---|---|---|
| All Events | 100,000 | #### | 6,350 | 1,336 | 440 | 335 | 410 | 93.2 | 8 | 7 | 577 | 449 |
| P1 | 99,708 | 99.7 | 6,364 | 1,337 | 440 | 336 | 410 | 93.2 | 8 | 7 | 575 | 449 |
| P2 | 88,071 | 88.1 | 5,604 | 1,174 | 387 | 311 | 357 | 92.4 | 7 | 7 | 507 | 419 |
| P3 | 86,310 | 86.3 | 5,473 | 1,154 | 379 | 307 | 350 | 92.3 | 7 | 6 | 497 | 415 |
| P4 | 99,786 | 99.8 | 6,358 | 1,337 | 439 | 336 | 378 | 86.1 | 8 | 7 | 576 | 449 |

**Figure S21** – A snapshot of flow cytometry experiment with GFP tagged strains

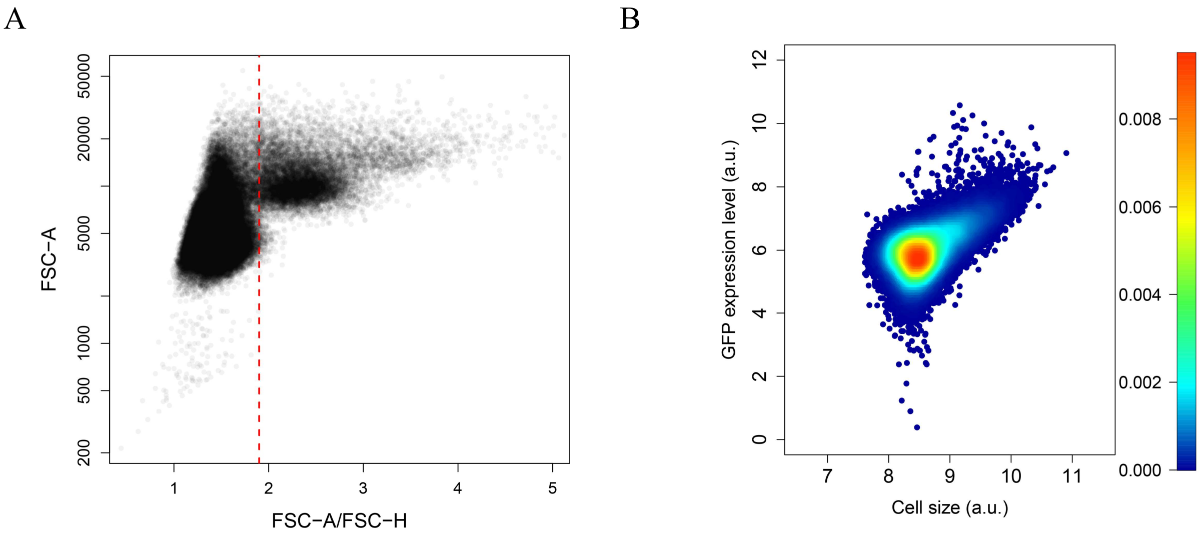


**Figure S22 – Processing of flow cytometry data for quantification of fluorescence signal from Rh123 staining, GFP tagged strains and Nile red staining (A)** FSC-A vs FSC-A/FSC-H plot that was used for filtering out cell aggregates. A cutoff of 1.9 for FSC-A/FSC-H was chosen. Events with FSC-A/FSC-H ratio lower than the cutoff value were considered as single cells. **(B)** GFP expression vs cell size (FSC-A) for a ICL1-GFP construct in A1 line.

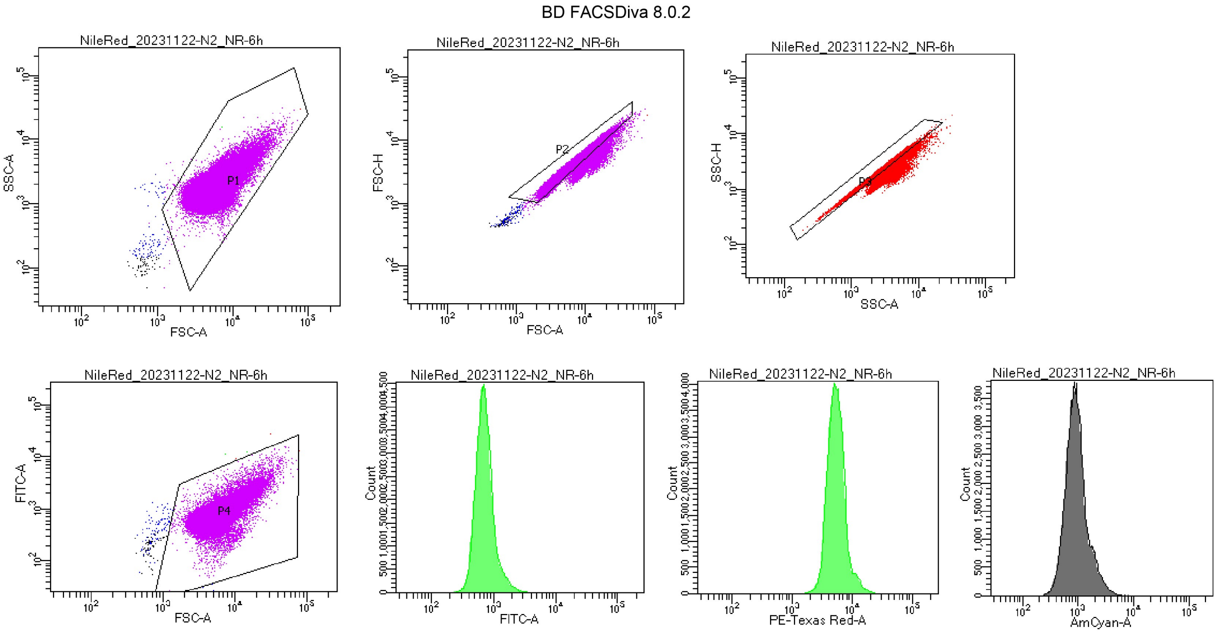


Tube Name: N2_NR-6h
GUID: b4d55ed2-2609-4344-83d3-4b2b658a46f5

| Population | #Events | %Parent | FSC-A Mean | SSC-A Mean | FITC-A Mean | FITC-A Median | FITC-A SD | FITC-A %CV | PE-Texas ... Mean | PE-Texas ... Median | AmCyan-A Mean | AmCyan-A Median |
|---|---|---|---|---|---|---|---|---|---|---|---|---|
| All Events | 100,000 | #### | 6,184 | 1,924 | 824 | 680 | 580 | 70.4 | 6,068 | 5,198 | 1,009 | 836 |
| P1 | 99,810 | 99.8 | 6,193 | 1,926 | 825 | 681 | 580 | 70.3 | 6,076 | 5,202 | 1,010 | 836 |
| P2 | 90,387 | 90.4 | 5,567 | 1,720 | 736 | 657 | 401 | 54.4 | 5,464 | 4,997 | 897 | 800 |
| P3 | 88,076 | 88.1 | 5,341 | 1,657 | 706 | 650 | 315 | 44.6 | 5,267 | 4,939 | 857 | 789 |
| P4 | 99,838 | 99.8 | 6,191 | 1,925 | 824 | 681 | 566 | 68.7 | 6,072 | 5,201 | 1,009 | 836 |

**Figure S23 –** A snapshot of flow cytometry experiment after Nile red staining

Supplementary tables

**Table S1 -** Primers for creating GFP-tagged fusion proteins of *PDC1*, *ICL1* and *SOD1* genes. The F2, R1 and CHK primers were taken from Howson *et al*., 2005.

| **Primer** | **Primer Sequence 5' to 3'** |
|---|---|
| PDC1_F2 | TGAACAAGCTAAGTTGACTGCTGCTACCAACGCTAAGCAAggtcgacggatccccgggtt |
| PDC1_R1 | TGCTTATAAAACTTTAACTAATAATTAGAGATTAAATCGCtcgatgaattcgagctcgtt |
| ICL1_F2 | TGTCACAGAAGATCAATTCAAAGAAAATGGCGTAAAGAAAggtcgacggatccccgggtt |
| ICL1_R1 | TGTCAGGAAATGCCGGCAGTTCTAATGGTTAATCCTTGTCtcgatgaattcgagctcgtt |
| SOD1_F2 | CGGTCCAAGACCAGCCTGTGGTGTCATTGGTCTAACCAACggtcgacggatccccgggtt |
| SOD1_R1 | ACTTACATACGGTTTTTATTCAAGTATATTATCATTAACAtcgatgaattcgagctcgtt |
| PDC1_F2CHK | CTGCTGTCCCAGCTTCTACC |
| GFP_RevCHK | GCATCACCTTCACCCTCTCC |
| ICL1_F2CHK | CCCACAGAGAAGCCAAGAAG |
| SOD1_F2CHK | CGGAAAAACAGGCAAGAAAG |
